# The Role of Biopsy Position and Tumor-Associated Macrophages for Predictions on Recurrence of Malignant Gliomas: An In Silico Study

**DOI:** 10.1101/2024.06.25.600613

**Authors:** Pejman Shojaee, Edwin Weinholtz, Nadine S. Schaadt, Haralampos Hatzikirou

## Abstract

Predicting the biological behavior and time to recurrence (TTR) of high-grade diffuse gliomas (HGG) after the maximum safe neurosurgical resection and combined radiation and chemotherapy plays a pivotal role in planning the clinical follow-up, the choice of potentially necessary second-line treatment, and the quality of life of patients faced with the diagnosis of a malignant brain tumor. The current standard-of-care (SoC) for HGG includes follow-up neuroradiological imaging to detect recurrence as early as possible and several clinical, neuropathological, and radiological prognostic factors with limited accuracy toward predicting TTR. Herein, using an in-silico analysis, we aim to improve predictive power towards TTR considering the role of (i) prognostically relevant information available by diagnostics used in current SoC, (ii) advanced image-based information that is currently not part of the standard diagnostic workup, such as interface of tumor and normal tissue (edge) features and quantitative data specific for the position of biopsies within the tumor, and (iii) information on tumor-associated macrophages. In particular, we introduce a state-of-the-art spatio-temporal model of tumor-immune interactions, emphasizing the interplay between macrophages and glioma cells. This model serves as a synthetic reality for assessing the predictive value of various features. We generate a cohort of virtual patients based on our mathematical model. Each patient’s dataset includes simulated T1 and FLAIR MRI volumes, and simulated results on macrophage density and proliferative activity either in a specified part of the tumor, namely tumor core or edge (”localized”), or unspecified (”non-localized”). We impose different levels of noise to enhance the realism of our synthetic data. Our findings reveal that macrophage density at the tumor edge contributes to a high predictive value of feature importance for the selected regression model. Moreover, there is a lower MSE and higher *R*^2^ for the ”localized” biopsy in prediction accuracy toward recurrence post-resection compared with ”non-localized” specimens. In conclusion, the results show us that localized biopsies can bring more information about the tumor behavior, especially at the interface of tumor and normal tissue (Edge).

## 1 Introduction

Diffuse high-grade diffuse gliomas (HGG) include the most common types of primary malignant brain tumors in adults namely glioblastoma multiforme (GBM), astrocytoma grade 3/grade 4 (A°4/A°3, and oligodendroglioma grade 3 (O°3) [1]. Glioblastoma is associated with the most unfavorable prognosis and poor significant therapeutic advances in the past few decades [2], and A°4/A°3 with better prognosis but a similar need for aggressive therapy and almost inevitable recurrence [3, 4, 5, 6]. HGG shares their potential for diffuse invasion of adjacent CNS structures and their phenotypical plasticity. In contrast, phenotypic plasticity is recognized as one of the hallmarks of cancer[7] and can be associated with tissue invasion in many types of cancer [8, 9, 10] the specific feature of glioma cells is ability to undergo transition to migratory phenotypes, in response to cues from the tumor microenvironment (TME). Due to the special anatomy of the brain and processes allowing glioma cells to acquire features of immature glia, this enables malignant cells to migrate through the brain extracellular matrix (ECM) towards distant sites in the CNS. The phenotypic shift is mainly due to the TME signals and local cell density which triggers the glioma cells toward migration. These infiltrating glioma cells are among the major reasons that make gliomas difficult to treat and are considered to be the starting point for recurrence. After the first line of treatment including maximal safe resection [11], this small population of remaining surviving infiltrating glioma cells will almost inevitably start to ”grow”, or ”go”. This leads to the need for tight post-operative follow-up to detect and treat recurrence, the major concern of second-line treatment and clinical decision-making in HGG[12, 13].

Predicting the recurrence time for glioblastoma multiforme (GBM) presents significant challenges due to limitations in current medical follow-up and diagnostic technologies. Although Magnetic Resonance Imaging (MRI) and biopsies are routinely employed following tumor detection, these methods have inherent limitations that complicate the accurate prediction of relapse. The extent of tumor resection is often imprecisely known, which is crucial for predicting recurrence since residual tumor cells can drive relapse. Additionally, the spatial variability in biopsy locations can lead to sampling errors, as the biopsies may not accurately represent the tumor’s heterogeneity. This variability impedes the development of reliable biomarkers for predicting recurrence times, thus obstructing the formulation of an effective, personalized post-treatment monitoring strategy for GBM patients. These challenges underscore the need for advancements in both diagnostic techniques and post-treatment surveillance protocols to improve prognostic assessments and therapeutic outcomes in GBM care [14, 15, 16].

Here, we focus on a specific type of plasticity between proliferative and invasive cell phenotypes, the so-called Go-or-Grow mechanism, which has an important role in tumor progression and recurrence [17, 18, 13, 19]. In particular, proliferative cells have a lower tendency to infiltrate other regions of the brain, and migratory cells have a lower proliferation rate. Fig. 1 shows examples of proliferative and infiltrative types of tumor types which have short or long-distance invasive edges. Some studies linked the Go-or-Grow behavior to the effects of TME stimuli [20, 21, 22, 23, 19] as well as immune system engagement in the TME [24, 25, 26, 27, 28, 29]. In this study, we consider the effects of oxygen and macrophages as the main role players in the TME to interact with glioma cells. Macrophages are the most abundant immune cells in the TME with up to (30%–50%) of the tumor mass in the TME [24, 30, 29]. Tumor-associated macrophages (TAMs), which include both brain-resident microglia and macrophages originating from bone marrow, are attracted to the tumor site by various cytokines and chemokines. These signaling molecules, including CXC motif chemokine ligand 16 (CXCL16), CC motif chemokine ligand 2 (CCL2), transforming growth factor beta (TGF-*β*), and interleukin 33 (IL-33), play a crucial role in their recruitment [31]. TAMs are a heterogeneous population not only upon their origin but also on their functionality in TME. TAMs are classically classified into binary phenotypes as a pro-inflammatory M1 which is characterized by the pro-inflammatory cytokines such as IL-1*β* and TNF plus the activation of immune receptors TLR2/4 and M2 phenotype, known for its anti-inflammatory properties, is characterized by the production of ARG1, IL-10, and IL-4 [30, 32]. TAMs are predominantly found in the hypoxic areas of tumors. Their presence is likely due to the combined influences of various chemoattractants, which draw the macrophages in, and the limited mobility within these low-oxygen environments, which causes the macrophages to accumulate. Once in these regions, the macrophages tend to transition into the M2 subtype, adopting functions that promote tumor growth. The combination of the M2 phenotype of TAMs and intratumoral hypoxia contributes to enhanced tumor malignancy and significantly reduces the effectiveness of immunotherapy [33, 34, 35, 36].

**Fig. 1.**
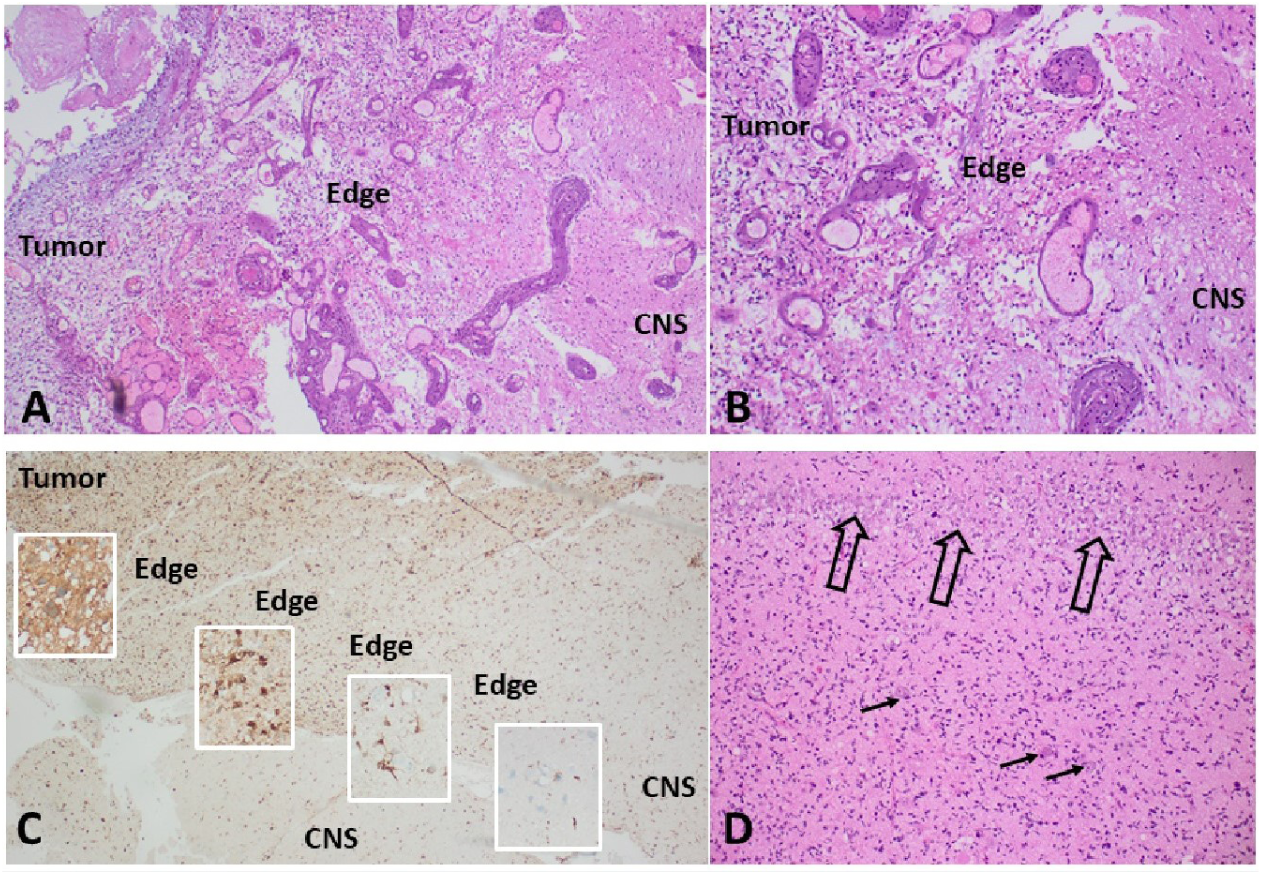
A and B: Examples of short, almost “nodular” invasion front in an IDH wild type glioblastoma (WHO grade 4), demonstrating a short distance from the solid tumor to the leading edge and pre-existing CNS tissue. C and D: Examples of long-distance invasive edges in a diffusely infiltrating malignant glioma (IDH1 mutated astrocytoma, WHO grade 4). C: Immunohistochemistry for R132H mutated IDH1, highlighting the tumor cells in brown (DAB) staining. D: Detail from the area in the middle of the long invasive edge, showing single pre-existing neurons of the temporal cortex (arrows) and the dense band of hippocampal pyramidal cells (open arrows), both diffusely infiltrated by tumor cells. (A, B, D: Hematoxylin/Eosin staining)

To dissect the aforementioned complexity, researchers developed appropriate mathematical models. Such models enable biologists and clinicians to quantitively understand and forecast tumor dynamics [37]. They can be valuable not only in general biological research but also in precision medicine by bridging the knowledge gaps between TME heterogeneity and cellular plasticity. In the context of mathematical biology, several models including discrete, continuous, and hybrid have been deployed to describe the tumor-immune interactions based on the plasticity of the cells [38, 29]. Several review papers provide an overview of the various approaches for mathematical models in this domain [29, 39, 40, 41, 42].

In this study, we pose the following questions:

**Question 1**. Which role do TAMs play in aiding the invasive edge of tumor cells to facilitate recurrence?

**Question 2**. Can we delineate the complex dynamics that emerge by the interactions of two phenotypically plastic populations, namely the glioma cells and TAMs in the GBM microenvironment?

**Question 3**. How can we use the model to explore the prognostic power of localized and non-localized biopsies through an *in-silico* base?

To answer these questions, we developed a state-of-the-art mathematical model based on a previous study by Alfonso *et al.*, 2016 [38, 37]. This spatiotemporal PDE model involves TAMs interacting with glioma cells and modulated by the oxygen gradients. In turn, we analyzed the model inside a parameter space (proliferation and infiltration rate) for the two clinical outputs, namely infiltration width and tumor size. Then, we analyzed the parameters of the system with a global time-dependent sensitivity analysis to identify the most effective parameter throughout the time (Q1 and Q2). In the second part, we generated a virtual cohort and included a resection for each patient-specific tumor detection and continued to simulate after the resection to obtain the IW after 3 and 12 months post-resection for each virtual patient. Then, we divided the virtual cohort into datasets reflecting the predominantly diffuse/infiltrative behavior as opposed to the predominantly nodular growth and analyzed the respective prediction values which is the predicted IW via a selected regression model (Q3). It is important to note that the term ”nodular” is typically reserved for certain types of nodular growth that resemble clusters of grapes, unlike the less well-demarcated glioma invasive front. In our context, while we use the term ”nodular”, we acknowledge that all high-grade gliomas (HGG) exhibit diffuse invasive infiltration of the central nervous system (CNS). Here, our understanding of ”nodular” refers to a ”narrow invasive edge” as opposed to more long-distance diffuse infiltration. Fig. 2 shows an overview of the whole procedure with more detail.

**Fig. 2.**
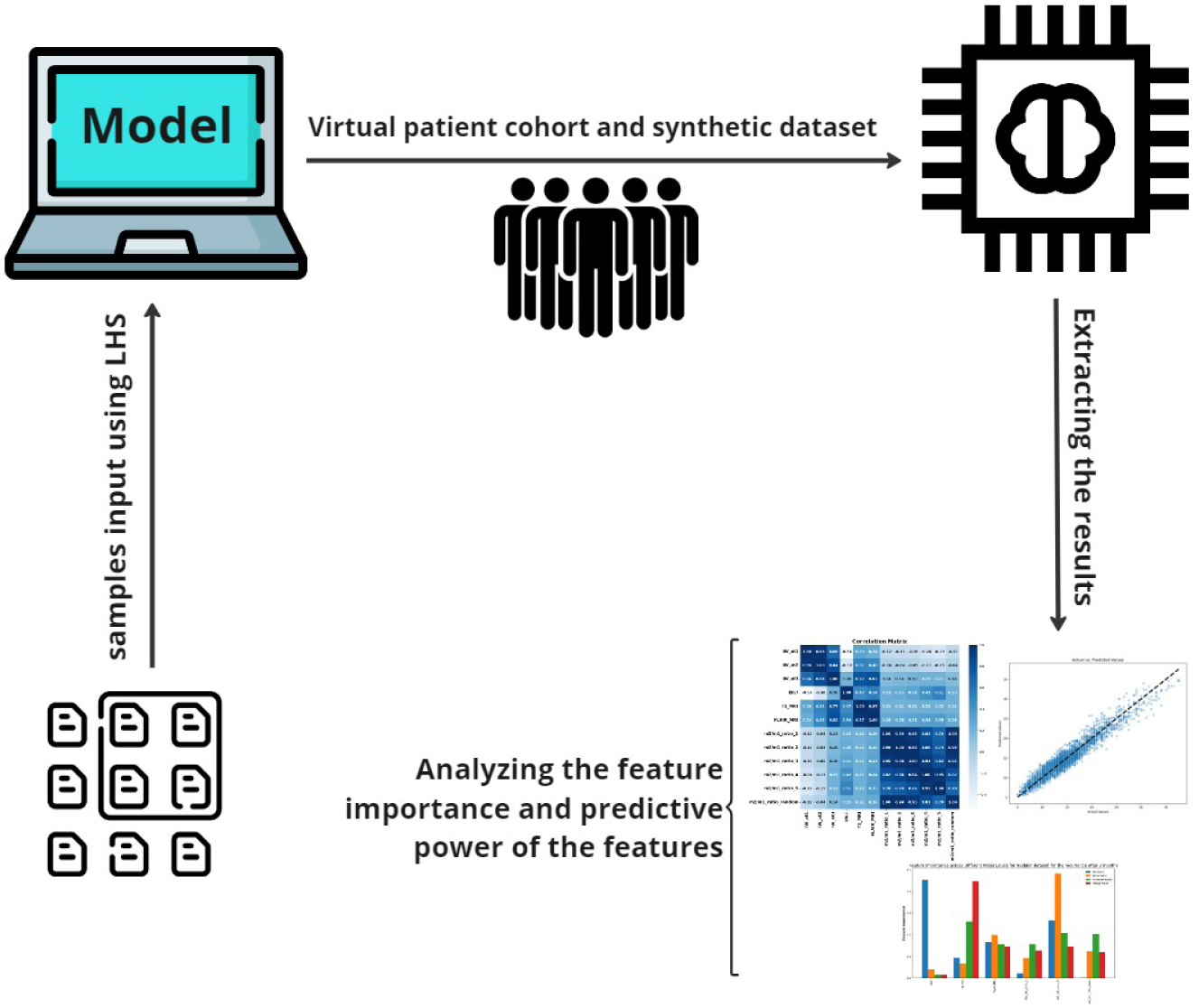
Overview of the Study Procedure. Our study started with the generation of a comprehensive list of 20,000 parameters utilizing the Latin Hypercube Sampling (LHS) method. These parameters were then integrated into our mathematical model to simulate the infiltration widths after the resection for 3 to 12 months. The resultant data facilitated the creation of a synthetic dataset, emulating a patient cohort. This dataset served as the foundation for our subsequent analysis, employing machine learning techniques to unravel the predictive capabilities of the model and to discern the significance of various features within it (Icon used from Flaticon.com).

## 2 Materials and methods

### 2.1 Mathematical modeling of the interactions between glioma and macrophages

In this study, macrophages were added to a system of PDE equations, playing an active role in interacting with glioma cells both in inhibiting and activating glioma growth and infiltration. The basic model was developed previously by considering tumor vasculature and oxygen concentration [38, 37]. In this work, we assume tumor vasculature as an implicit parameter that influences the influx of TAMs and oxygen transport, and for simplicity, we assume it as a constant value. Fig. 3 shows a schematic of the model’s variables interactions and how macrophages impacts on the dynamic of gliomas with the main assumptions. The variables are *ρ_m_*(*x, t*) as the glioma infiltrative cells (Go), *ρ_p_*(*x, t*) as the glioma proliferative cells (Grow), *n*(*x, t*) refers to the amount of oxygen as the nutrient factor, *m*_1_(*x, t*) refers to the anti-tumor TAMs, and *m*_2_(*x, t*) refers to the pro-tumor TAMs; where (*x, t*) ∈ R*^d^* × R_+_ and *d* is the dimension of the system.

**Fig. 3.**
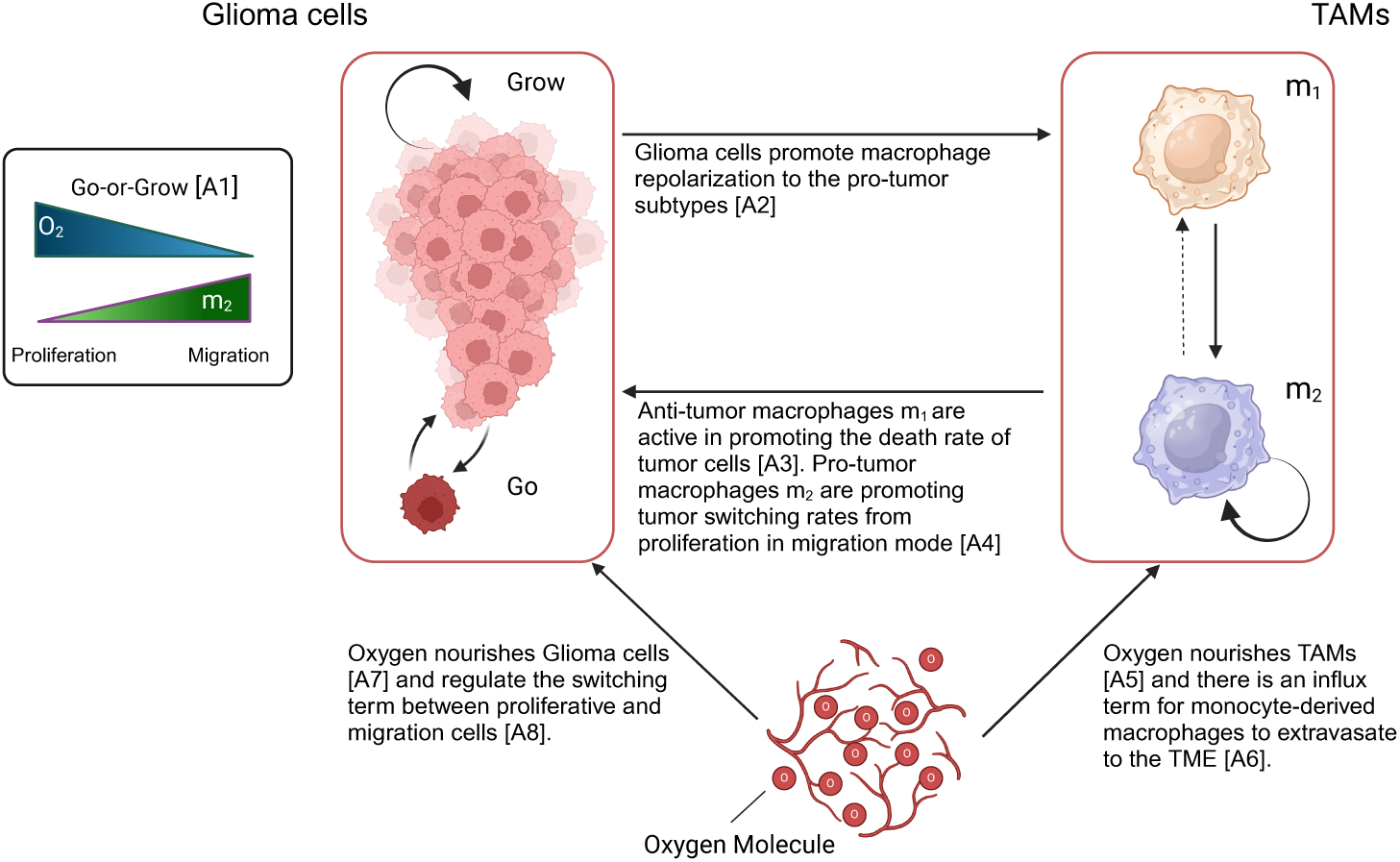
Scheme of the interplay between the tumor and TAMs cells by considering the role of oxygen in a spatial form of the model. The scheme shows a holistic view of the interactions and switching terms both for glioma and TAMs cells in the TME. (Created with BioRender.com)

The main assumptions are as follows:

A1 Glioma cells can assume two different phenotypes, namely proliferative or migratory (infiltrative) cells which depend on the TME stimulants. High (low) oxygen availability and low (high) pro-tumor TAMs induce proliferative (migratory) phenotypes [38, 37, 17, 43, 19, 44].

A2 Glioma cells significantly influence the macrophage polarization in the TME employing key cytokines like IL-4/13 and TGF-*β* playing a pivotal role [45, 30].

A3 M1-like macrophages are characterized by features working against tumor progression, primarily due to their intrinsic capabilities of phagocytosis and the generation of enhanced anti-tumor inflammatory responses [46, 47].

A4 Pro-tumor (”M2-like”) TAMs tend to increase the tumor invasive behavior. Therefore, we assume that M2-like macrophages are promoting the phenotypic switching rate from proliferation to the migratory glioma cells [44].

A5 and A7 Oxygen nourishes Glioma and TAMs cells in the TME [48, 49, 50].

A6 Monocyte-derived macrophages enter the CNS via the tumor vasculature [27].

A8 Oxygen concentration plays a specific role in changing the glioma phenotypes [48, 49].

### 2.2 Glioma cell dynamics

We introduce the density of migrating and proliferating glioma cells, denoted by *ρ_m_* and *ρ_p_*, respectively. The switching term from proliferation to migration depends on the pro-tumor macrophages (*m*_2_) and oxygen concentration (*n*).

The equations for the two distinct glioma phenotypes are as follows:

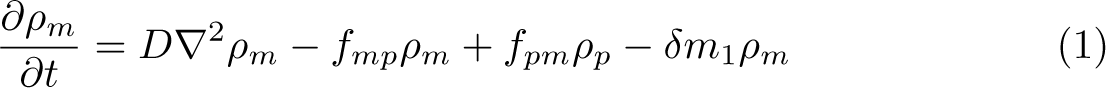

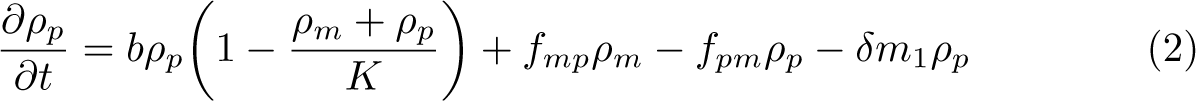

where *f_mp_ρ_m_*and *f_pm_ρ_p_*are the switching terms between the two cell phenotypes. The switching parameters (*f_mp_* and *f_pm_*) are dependent on the oxygen and M2-like macrophages. The system could be reduced to a single equation with a density for both glioma cell phenotypes. This assumption appears reasonable, as intracellular mechanisms like signaling pathways that control the phenotypic transition function on significantly shorter time scales than those of cell migration and proliferation [38, 37]. Consequently, it is presumed that the switching of phenotypes occurs more rapidly than cell division and movement, enabling the expression of *ρ_m_* and *ρ_p_*concerning *ρ*. We can express a *ρ* = *ρ_m_* + *ρ_p_* and *f_pm_ρ_p_*= *f_mp_ρ_m_* as follows:

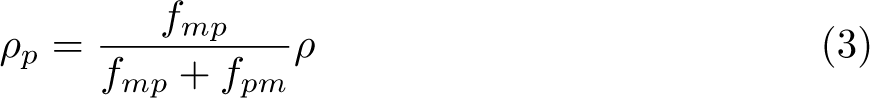

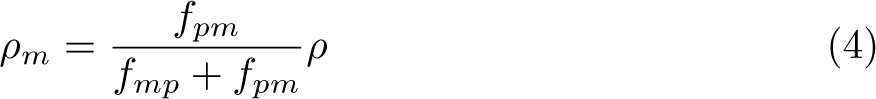

Thus, we can express the glioma density as follows:

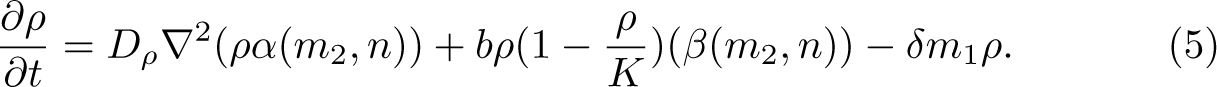

*α*(*m*_2_*, n*) and *β*(*m*_2_*, n*) were defined as:

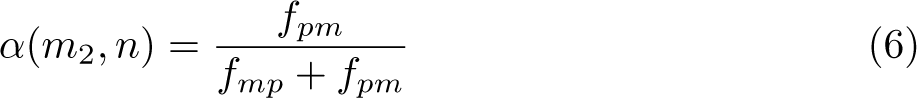

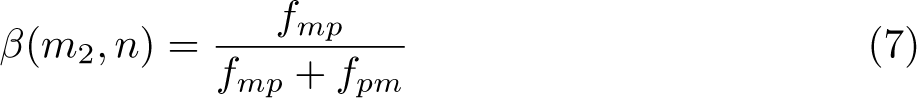

#### 2.2.1 Glioma phenotypic switching term

As we briefly described in Fig. 3, glioma cells can change their phenotype into migration or proliferation due to the oxygen and effect of TAMs. Based on prior studies, we incorporated a linear function of oxygen into the phenotypic switching model [38, 37]. Whereas, we assumed *m*_2_*/K_m_* for the TAMs to represent the effect of pro-tumor macrophages on the glioma cells. TAMs release factors (IL-6, IL-1*β*, EGF, TGF-*β*, STI1, and PTN) that promote tumor progression [30]. This assumption implies the amount of pro-tumor macrophages in the TME. The function can be tailored to fit experimental data or specific biological scenarios. In this sense, we defined the switching term as below:

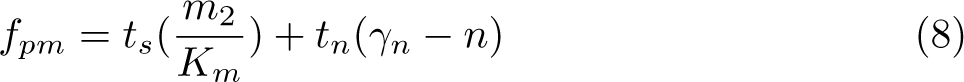

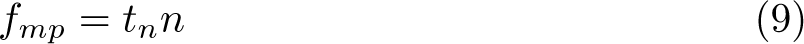

*γ_n_*is the small regularization term for the possible transition when oxygen is zero. Fig. 4 shows the 2D plot behavior of 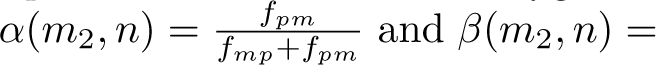 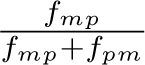 for the different values of *m*_2_ and *n*.

**Fig. 4.**
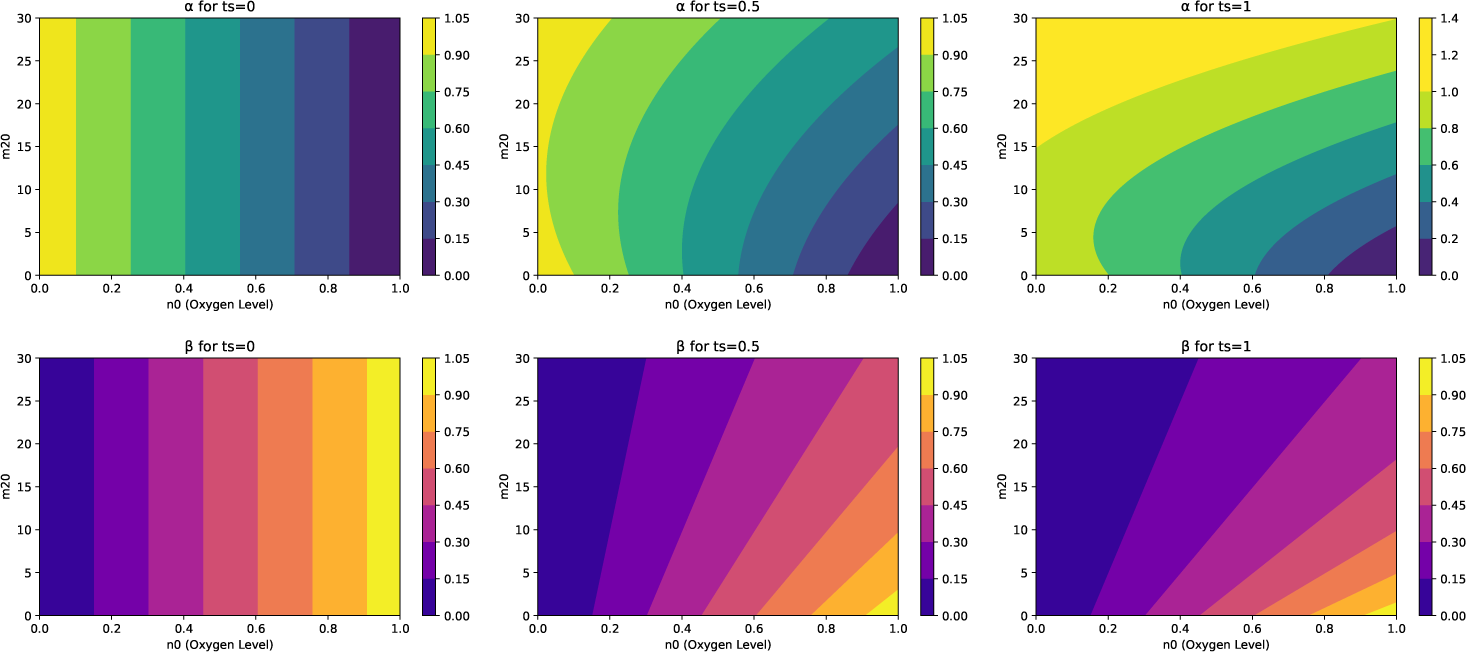
Contour plots showcasing the dependence of *α* and *β* values in relation to oxygen levels (*n*) and TAMs (*m*_2_) at three different *t*_s_ (ts=0, 0.5, 1). These plots reveal the dependency of switching terms, indicative of the migratory cell transition rate within the Tumor Microenvironment (TME), on the interplay between environmental oxygen and the M2-like parameter, emphasizing the complexity of cellular behavior in response to the TME.

Then, the transition terms for *α*(*m*_2_*, n*) and *β*(*m*_2_*, n*) become:

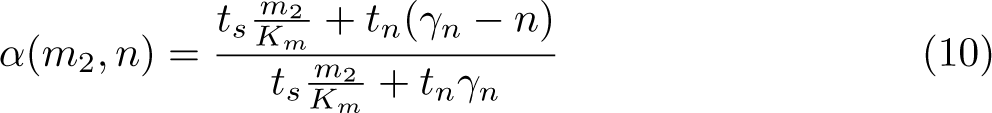

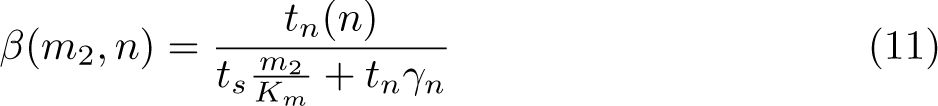

The terms *α*(*m*_2_*, n*) and *β*(*m*_2_*, n*) encapsulate the central role of oxygen and TAMs in the TME, influencing glioma plasticity and regulating diffusion and proliferation rates of glioma cells based on the availability of oxygen and TAMs. The nonlinearity of these terms arises from the division of linear components, resulting in a ratio that does not adhere to linearity concerning either variable. Consequently, *α*(*m*_2_*, n*) and *β*(*m*_2_*, n*) represent nonlinear functions of *m*_2_ and *n*. We hypothesize that TAMs facilitate tumor infiltration, while oxygen levels simulate normoxic and hypoxic regions where the tumor tends to grow or infiltrate, respectively.

### 2.3 TAMs dynamics in the TME

To model the TAMs in the spatiotemporal domain, we simplified the known broad spectrum of macrophage polarization and simplistically assumed two major populations: pro-inflammatory (*m*_1_) and anti-inflammatory (*m*_2_) TAMs. In this simplified concept, neglecting the pre-existent macrophage-like population of microglia, the *m*_1_ macrophages have an influx based on the amount of tumor vasculature. We also added a diffusion term to incorporate the infiltration of TAMs in the TME. The role of glioma cells influences the switching term between pro-tumor and anti-tumor TAMs. Glioma cells release chemoattractant molecules (Periostin, SSP1, GM-CSF, SDF-1, ATP, GDNF, CX3CL1, LOX, HGF/SF, MCP-1, MCP-3, CSF-1, Versican, and IL-33) [30] that recruit and reprogram TAMs to facilitate their tumor progression. The equations for *m*_1_ and *m*_2_ are as follows:

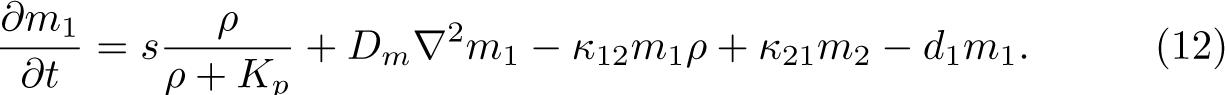

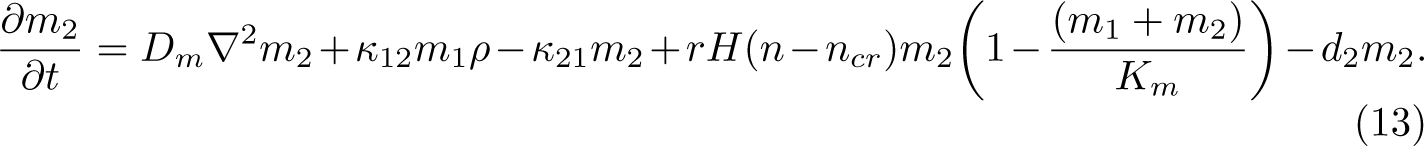

where *D_m_* is the diffusion term for macrophages and *κ*_12_, *κ*_21_ are the switching rates between two extreme macrophage phenotypes. *K_p_* is the half-saturated rate of glioma cells for macrophage recruitment, and *S* is the recruitment coefficient. 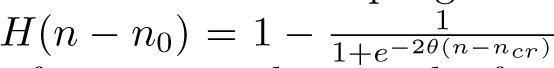 is the heavy-side function that shows the effect of oxygen on the growth of pro-tumor TAMs. This term models the hypoxic and normoxic regions of the TME, where pro-tumor macrophages are prone to accumulate in hypoxic areas. The accumulation of TAMs in hypoxic zones within tumors is often linked to a negative outcome in clinical settings [35, 36]. These macrophages are drawn to these low-oxygen areas by following the trail of certain chemical attractants. Once there, TAMs contribute to tumor growth and progression through various mechanisms. They promote the formation of new blood and lymph vessels (angiogenesis and lymphangiogenesis), assist in the spread of tumor cells (metastasis), and suppress immune responses, thereby aiding the tumor’s survival and expansion [35, 36, 51].

### 2.4 Oxygen concentration dynamics

Oxygen is delivered to the brain tissue by the vascular network in the brain and by the functional blood vessels through the tumor bulk to be consumed by the cells such as glioma and TAMs in the TME. The oxygen transport in the brain is conducted by diffusion and convection [52]. In this work, we only consider the diffusion form of oxygen transport throughout the TME. Oxygen is delivered by the functional vasculature and the supply rate is proportional to the difference between normal oxygen concentration in the brain *n*_0_ and the TME.

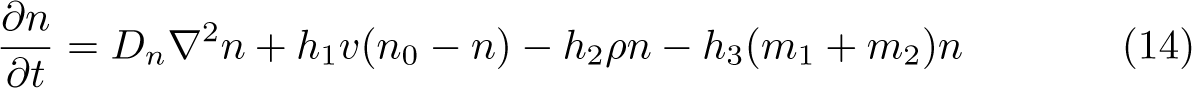

where *D_n_* is the oxygen diffusion coefficient, *h*_1_ is the permeability coefficient of functional vasculature, *v* is the constant parameter for the vasculature, we assume a constant value which is uniform through the space, *h*_2_ is the oxygen consumption range by the glioma cells, and *h*_3_ is the TAMs oxygen consumption rate.

#### 2.4.1 Model parameterisation

The values for parameters used in the model’s simulations are primarily sourced from existing literature. When specific data is not available, estimates were made to closely align with physiological conditions, guided by relevant physical and biological reasoning. In this case, we estimate the values for TAMs diffusion rates *D_m_*, TAMs consumption rate *h*_3_, and regularization terms *t_s_* and *t_n_* based on the closest relevant physiological conditions. For the sensitivity analysis, a wide range of values is chosen based on the interesting parameters to show their sensitivity and correlation to the observable outputs. Table 1 depicts the parameter values used in this study.

**Table 1.** Parameter values and specific ranges in the model simulations

| Parameter | Value | Unit | Description (source) |
| --- | --- | --- | --- |
| $D$ | $[2.73 \times 10^{-3}, 2.73 \times 10^{-1}]$ | $mm^2 day^{-1}$ | Diffusion rate of glioma cells [53, 54, 52] |
| $b$ | $[2.73 \times 10^{-4}, 2.73 \times 10^{-2}]$ | $day^{-1}$ | Glioma cell proliferation rate [53, 54, 52] |
| $\delta$ | $2 \times 10^{-4}$ | $day(cells)^{-1}$ | Glioma cell death rate induced by macrophages |
| $K$ | $10^2$ | $cells(mm)^{-1}$ | Tumor carrying capacity [55, 56] |
| $K_m$ | $10^2 \times 25\%$ | $cells(mm)^{-1}$ | macrophage carrying capacity [30] |
| $t_s$ | $[0, 0.5]$ | - | $m_1$ transition rate (Estimated) |
| $t_n$ | $[0, 0.5]$ | $nmol(day)^{-1}$ | nutrient transition rate (Estimated) |
| $d_1$ | 0.87 | $day^{-1}$ | $m_1$ death rate [57] |
| $d_2$ | 0.1 | $day^{-1}$ | $m_2$ death rate [57] |
| $r$ | $[0.487, 0.88]$ | $day^{-1}$ | Macrophage division rate [57] |
| $n_0$ | 1 | $nmol(day)^{-1}$ | Physiological oxygen concentration [58, 59] |
| $n_{cr}$ | 0.25 | $nmol(day)^{-1}$ | Critical oxygen concentration |
| $K_p$ | $0.5 \times K$ | - | saturation rate (Estimated) |
| $D_m$ | $[1 \times 10^{-3}, 1 \times 10^{-1}]$ | $mm^2 day^{-1}$ | Macrophage diffusion rate (Estimated) |
| $D_n$ | $1.51 \times 10^2$ | $mm^2 day^{-1}$ | Oxygen diffusion rate [60, 61, 62] |
| $h_1$ | $3.37 \times 10^{-1}$ | $day^{-1}$ | Oxygen supply rate [63, 64, 65] |
| $h_2$ | $[5.73 \times 10^{-3}, 1.14 \times 10^{-1}]$ | $mm(cell \cdot day)^{-1}$ | Glioma cell oxygen consumption rate [66, 67] |
| $h_3$ | $[1 \times 10^{-3}, 1 \times 10^{-1}]$ | $mm(cell \cdot day)^{-1}$ | Oxygen consumption rate by macrophages (Estimated) |
| $v$ | 1 | $cells(mm)^{-1}$ | Vasculature coefficient |
| $\kappa_{1,2}$ | $[10^{-5}, 10^{-2}]$ | $(day \cdot cells)^{-1}$ | M1 to M2 switch rate [57] |
| $\kappa_{2,1}$ | $[10^{-10}, 10^{-9}]$ | $(day \cdot cells)^{-1}$ | M2 to M1 switch rate [57] |
| $S$ | $[0.1, 0.5] \times K_m$ | $cells(mm)^{-1}$ | Recruitment rate (estimated) |
| $\theta$ | 10 | - | Slope parameter of the continuous Heaviside function $H$ |
| $\gamma_n$ | 1.01 | - | Regularization term |

### 2.5 The full model

The governing equations including a system of coupled partial differential equations which is used for the numerical simulation are as follows:

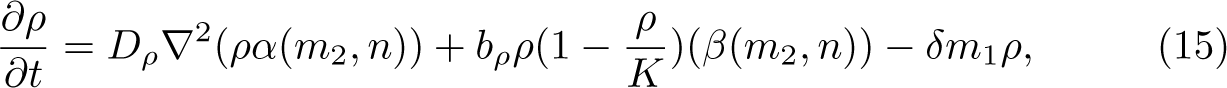

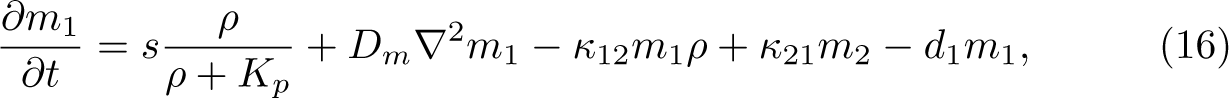

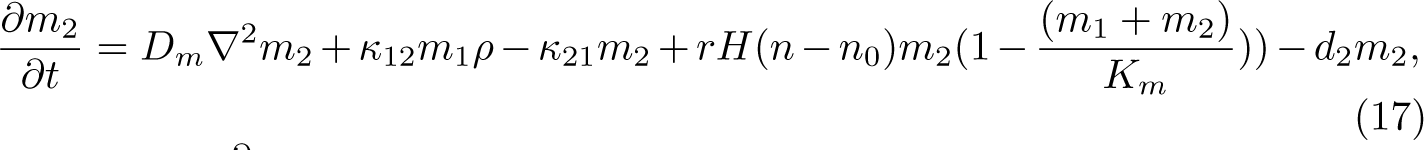

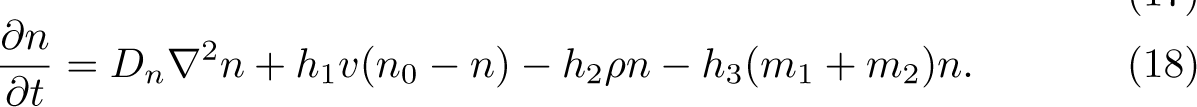

where oxygen and TAM-dependent switching terms are given by Eqs (8)–(9). We assume an initial density followed by:

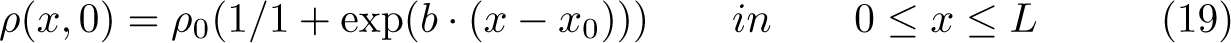

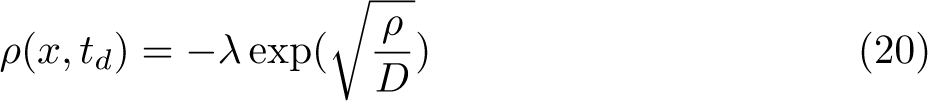

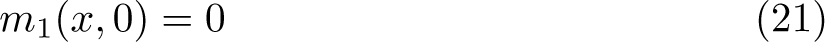

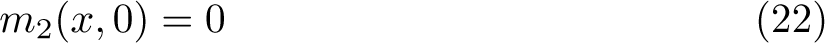

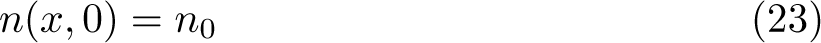

where the *m*_1_ and *m*_2_ have zero density at the beginning of calculations. *ρ*_0_ and *n* are the positive parameters and the density of glioma cells was distributed in a segment of length *x*_0_ in a sigmoidal form where *b >* 0 is a constant value. *L >* 0 is the length of a one-dimensional computational domain in *L* = 300*mm* which is discretized into a grid from 2.5×10*^−^*^3^ to 2.5×10*^−^*^2^ in a total simulation of about 3 years with a time-step of 30 days. *ρ*(*x, t_d_*) is the initial condition after the tumor resection where *λ* is the resection threshold [68]. To derive the exponential tail of a traveling wave solution of the model, we can neglect the non-linear term growth (*ρ*(*x*)*/K <<* 1) [68] and the role of *m*_1_ macrophages for inhibit<u>ing</u> the tumor growth rate. Therefore, we can obtain the value as 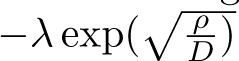 [68, 69].

Both grid network and time steps are selected properly to avoid numerical instability. Furthermore, we examine a standalone host tissue where all system dynamics are exclusively attributed to the interactions specified in Eqs. (16)-–(19). This premise leads to the implementation of fixed boundary conditions that prevent any flux.

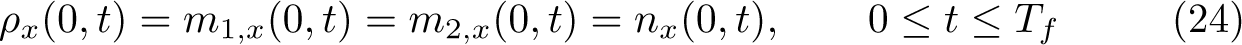

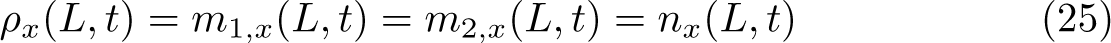

where *T_f_* is the end of simulation time which is three years in this study. In the no-flux boundary condition, no cell leaves through the boundaries.

### 2.6 Stability analysis

In the supplementary material, we provided the linear stability analysis for the glioma-free steady states of the diffusion-free model in dependence on these parameters. We found two such states which were non-negative. The trivial steady state [*ρ_p_, ρ_m_, m*_1_*, m*_2_*, n*] = [0, 0, 0, 0, 1] was associated with the healthy brain state since no monocyte-derived macrophages existed. We found this was a neutrally stable state with four negative and one close to zero (although positive) eigenvalue. For the special case of *t_s_*+ *t_n_* = 1, we were also able to infer the stability of another glioma-free steady state that existed for the hypoxic region in which we had *H* ≈ 1, where macrophages existed. This state was relevant in the case of a broken blood-brain barrier and eventually after tumor clearance. This state was stable only for a proliferation rate greater than a critical value *b_crit_*, whose value depended on a non-trivial combination of the other parameters. This could be understood as a local tumor-free state where macrophages were sufficient to clear the tumor.

#### 2.6.1 Model observable outputs

The model is used to produce two clinical outputs, namely tumor infiltration width (IW) and tumor size (TS). IW is the difference of distances from the core where the density of tumor mass is 80% and 2%, which is also considered as the tumor edge. TS was obtained by integrating the spatial profile of tumor density divided by its maximum value. These outputs represent the most critical parameters for estimating tumor malignancy and understanding the impact of various factors on them [38, 37, 53, 54, 70].

### 2.7 Machine learning analysis

To address Question 3 concerning the prognostic power of derived features, we have designed a methodology that evaluates various data availability scenarios, mirroring the post-surgical IW.

#### 2.7.1 Data preparation for generating clinical dataset

For generating the synthetic dataset, we adopt Latin Hypercube Sampling (LHS), the same strategy used in the sensitivity analysis of the preceding chapter but considering a uniform distribution for all the parameters. LHS is an effective approach for navigating the complex, multidimensional parameter spaces typical for mathematical biology models. It divides each parameter’s range into equal likelihood intervals and selects one sample from each, thus ensuring a broader distribution across the entire parameter space [71]. This method of stratification offers a more even coverage than simple random sampling from a uniform distribution, which often leads to sample clustering and inefficient exploration of the space. Utilizing LHS in creating the synthetic dataset results in a collection of parameter combinations that are more representative and varied, thereby enhancing the analysis of the model’s responses to different conditions.

In this section, we select key parameters that differ across the established clinical spectrum and our hypothesis to create 20,000 synthetic patients. These parameters serve as inputs for the model, from which we derive the concentrations of glioma, TAMs, and oxygen at a randomly determined detection time. The detection time varies for each patient, reflecting the unpredictable timing of MRI diagnoses of the tumor from initial growth. Upon determining the detection time for each synthetic patient, we set the tumor density to zero and treated the remaining tumor mass as the starting point for the new post-surgery condition. Fig. 5A illustrates the process of choosing the detection and prediction times and establishing the new baseline condition for each patient’s post-surgical profile. To compile the dataset, we simulated the acquisition of biopsies and MRI scans for patients. This method enables us to evaluate the predictive capacity of various features. We determine MRI data by utilizing 80% of the maximum tumor density for T1-weighted images and 16% for FLAIR images, which helps us estimate the tumor volume via these imaging techniques [54]. Additionally, we calculated the Ki67 biomarker, indicative of the patient-specific proliferation rate, using the formula proposed by Macklin *et al.* [72]:

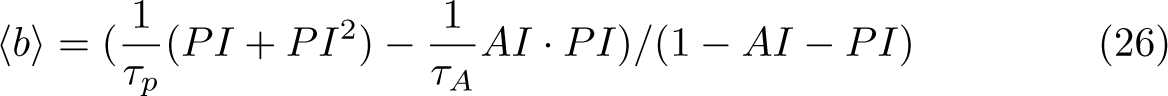

**Fig. 5.**
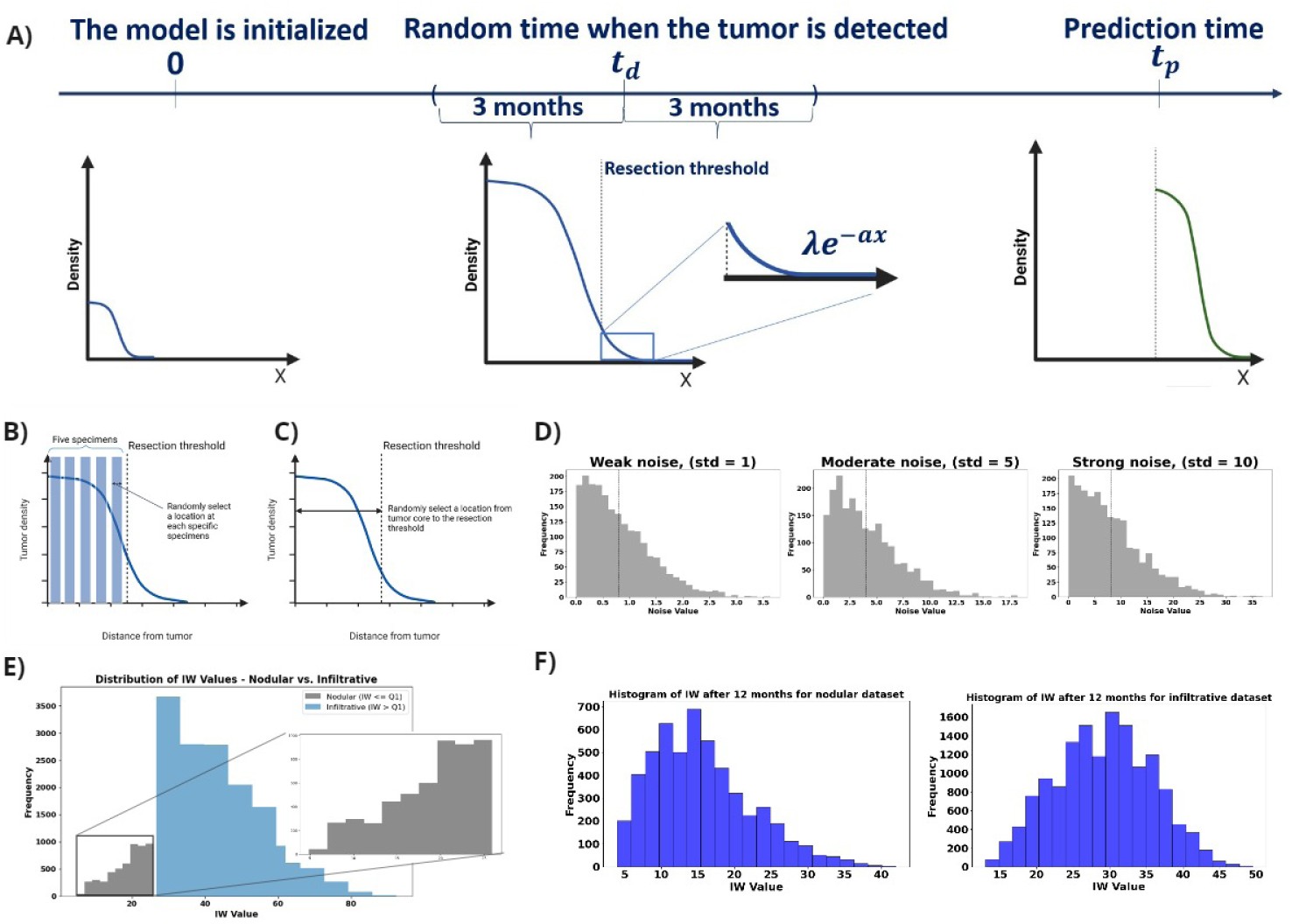
A) Initially, the model begins at *t* = 0, simulating tumor density, oxygen levels, and macrophage activity. For each simulated patient, a detection time is randomly assigned within a range of *±*3 months. Once the detection time for a patient is established, a hypothetical surgical resection is performed based on a predefined resection threshold. This process involves removing the resected area, leaving behind a truncated profile of tumor cells. These remaining cells are then considered the new starting point for further simulation, which continues until achieving our objective of generating clinical outcomes IW for 3, 6, and 12 months post-resection. In this schematic figure, B) we select five virtual ”specimens”, representing either biopsies or portions from the resected tumor, at the time of primary diagnosis. Then we extract the data including tumor cell and macrophage densities from these five simulated ”specimens”. C) we select random locations representing the full range from the tumor core to the resection margin. D) we impose Gaussian noise with three different standard deviations using a half-normal distribution for obtaining only positive values E) We divided the cohort into two subgroups: Nodular and Infiltrative based on the IW at the time of detection. This category allows us to investigate the predictive power of values across the range of formless aggressive to massively infiltrative tumors. F) The sample of histograms for the IW post-resection after 3 and 12 months for the infiltrative cohort. IWs are our target values for the prediction.

In this equation, PI represents the proliferation index, equated with the Ki67 values and AI denotes the apoptotic index, for which we fixed a value of 0.831%. The apoptosis duration *τ_A_*is assumed to be 8 hours, and the cell cycle time averages 25 hours, following the parameters provided by Macklin *et al.* and supported by additional sources [72, 73]. To collect the remaining biopsy data, which includes information on tumor cells and macrophages, we devise a strategy that segments the tumor from its core to its periphery into five distinct zones. We then select a random location within each zone to sample. At these chosen sites, we record the concentrations of tumor cells and macrophages. However, for the non-localized biopsy, we assume a random location from the tumor core until the resection threshold. Fig. 5B visually represents the process, depicting the biopsy samples extracted across the tumor’s profile. In the end, for the more realistic scenarios, we implement three different amounts of Gaussian noise with multiple standard deviations (Std =1, 5, 10) to make the biopsy data noisy. This procedure allows us to investigate the predictive power of biopsy data in different situations and quality. Fig. 5C shows the visualized noises which include weak, moderate, and strong noise based on the standard deviation.

#### 2.7.2 Evaluating predictive models for infiltration width in glioma subtypes

Understanding the difference between nodular and infiltrative gliomas is crucial for modeling tumor growth and response to therapies. Our approach began with categorizing the dataset into two distinct tumor subgroups: nodular and infiltrative (Fig. 5E). This classification was based on the infiltration width (IW) value observed at the time of tumor detection for each patient (Fig. 5F). Specifically, we designated tumors falling within the first quartile of IW values as nodular, while those exceeding this threshold were classified as infiltrative. Following this categorization, we embarked on a rigorous analysis to identify the most effective regression model for predicting IW at two critical post-resection intervals: 3 and 12 months. This selection process was informed by assessing the significance of various features within our dataset. Our methodology also incorporated two distinct biopsy techniques: localized and non-localized. it means that we take the values from the tumor core and the edge for the localized and a random pick-up from the range from the tumor core until the detection threshold. Moreover, we analyze the predictive power of each feature based on the metric values of MSE and *R*^2^ for the different case scenarios. Five case scenarios are introduced to predict the IW after 3 and 12 months. The scenarios are as follows:

I. [Ki67, T1 MRI, FLAIR MRI]
II. [Ki67, T1 MRI, FLAIR MRI, m2/m1 ratio]
III. [Ki67, T1 MRI, FLAIR MRI, m2/m1 at the core]
IV. [Ki67, T1 MRI, FLAIR MRI, m2/m1 at the edge]
V. [Ki67, T1 MRI, FLAIR MRI, m2/m1 at the core, m2/m1 at the edge]

where ’*m*_2_*/m*_1_ ratio’ refers to the non-localized biopsy, whereas ’*m*_2_*/m*_1_ at the core’ and ’*m*_2_*/m*_1_ at the edge’ refer to the biopsies from the tumor core and the closest region to the resection threshold. These scenarios depict the role of each feature, especially the role of *m*_2_ to *m*_1_ ratio for the localized and non-localized values. Fig. 6 shows the flowchart which depicts the holistic approach of this study. This procedure allows us to not only develop a mechanistic model but it can also be used as an in silico base to study the effect of each feature on the recurrence after the resection in virtual reality.

**Fig. 6.**
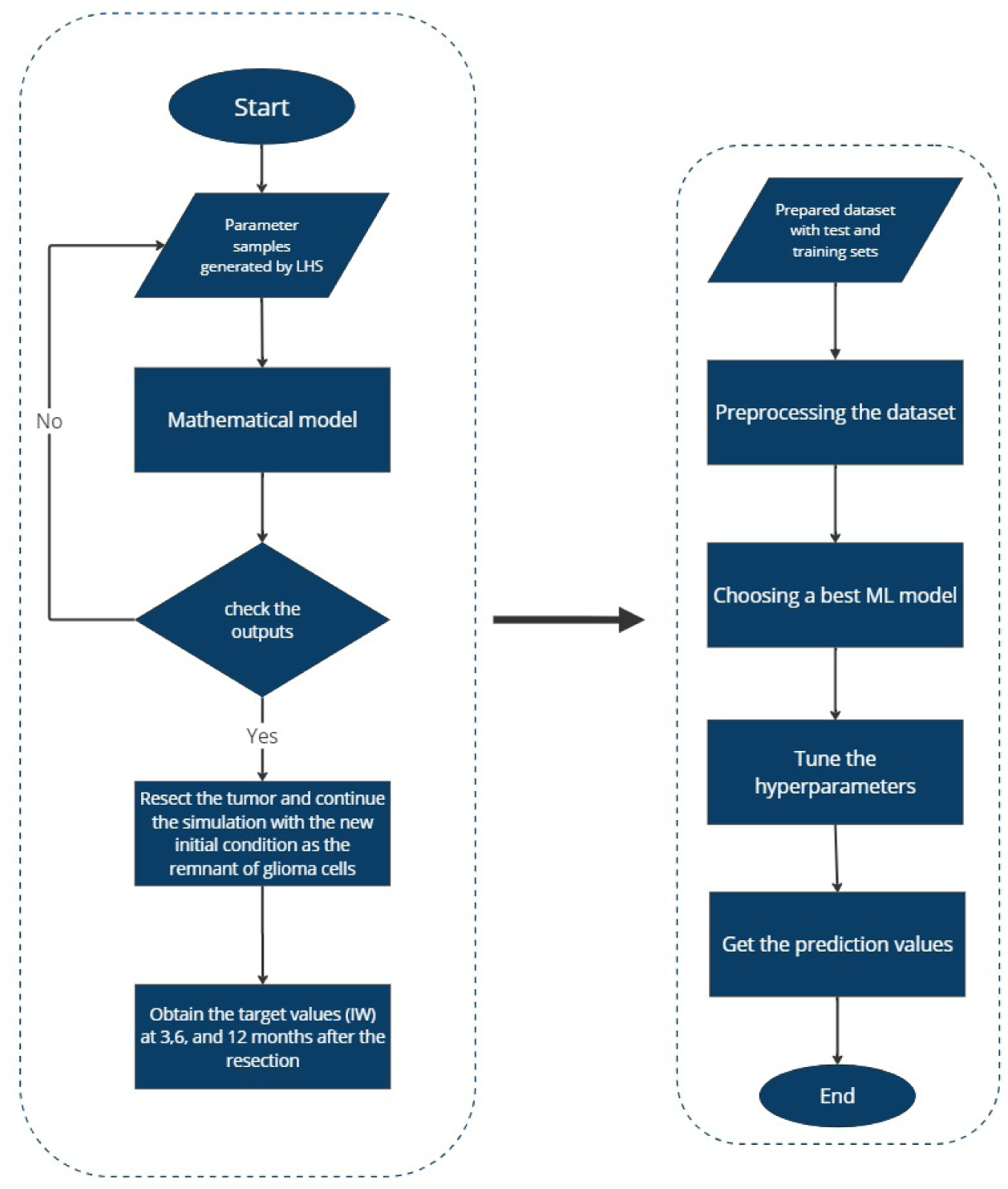
This flowchart illustrates a two-phase predictive modeling approach: first, a mathematical model simulates tumor dynamics based on parameters generated via Latin Hypercube Sampling (LHS) and post-surgical conditions; second, the simulated data informs a machine learning pipeline for the prediction of tumor progression at future time points to explore the prognostic power of features.

## 3 Results

We developed a model to address the key inquiries outlined in the Introduction. Our model encompasses tumor cells, TAMs, and oxygen levels, interlinked through a system of partial differential equations (PDEs). By analyzing the tumor’s behavior under the influence of TAMs and oxygen levels, we delved into understanding the dynamic interactions within the system. Initially, we evaluated the model’s performance against two clinical metrics: IW as infiltration width and Tumor Size (TS), illuminating the system’s behavior dynamics. Next, employing global sensitivity analysis, we probed the parameters to discern their impact on IW and TS outputs. Finally, leveraging the model as an in silico platform, we conducted simulations on a virtual cohort of 20,000 patients to predict IW levels at 3 and 12 months post-resection. This rigorous analysis not only unveiled the significance of localized biopsies but also provided insights into the predictive capabilities of individual parameters vis-à-vis target values.

### 3.1 The role of TAM and glioma cell interactions towards GBM growth and invasion

#### 3.1.1 Increasing glioma oxygen consumption rates increase the infiltrative behavior of tumor cells

We initiated our analysis by scrutinizing the impact of oxygen consumption rate on a parameter space (*D, b*). In Fig. 7, we presented a simulation map illustrating the IW across three distinct glioma oxygen consumption rates: *h*2 = [5.73 × 10*^−^*^4^, 5.73 × 10*^−^*^3^, 5.73 × 10*^−^*^2^]. Notably, simulations conducted under reduced oxygen consumption rates showed a prevalence of tumors characterized by decreased proliferation yet elevated infiltration rates, indicative of a propensity for invasive behavior. Moderate oxygen consumption rates, conversely, correlated with a marginal uptick in tumor proliferation rates, while the inclination toward invasiveness remains conspicuous. It was discerned that in normoxic conditions tumors tend towards nodular growth. However, escalating oxygen consumption rates corresponded with a pronounced shift towards invasive behavior across tumors, irrespective of their intrinsic proliferation rates. This phenomenon was attributed to hypoxia-induced cellular migration, fostering a tumor infiltrative pattern. Fig. 7 depicted the differential IW content in response to varying oxygen consumption rates, revealing subtle fluctuations in IW(II-I), in stark contrast to the pronounced disparities observed in IW(III-II). Specifically, a 1.96% increase in the maximum values of IW from IWI to IWII was noted, followed by a substantial 32.69% increase from IWII to IWIII. This latter observation underscored the pivotal influence of hypoxia in driving the transition of highly proliferative glioma cells toward a more aggressive and infiltrative phenotype.

**Fig. 7.**
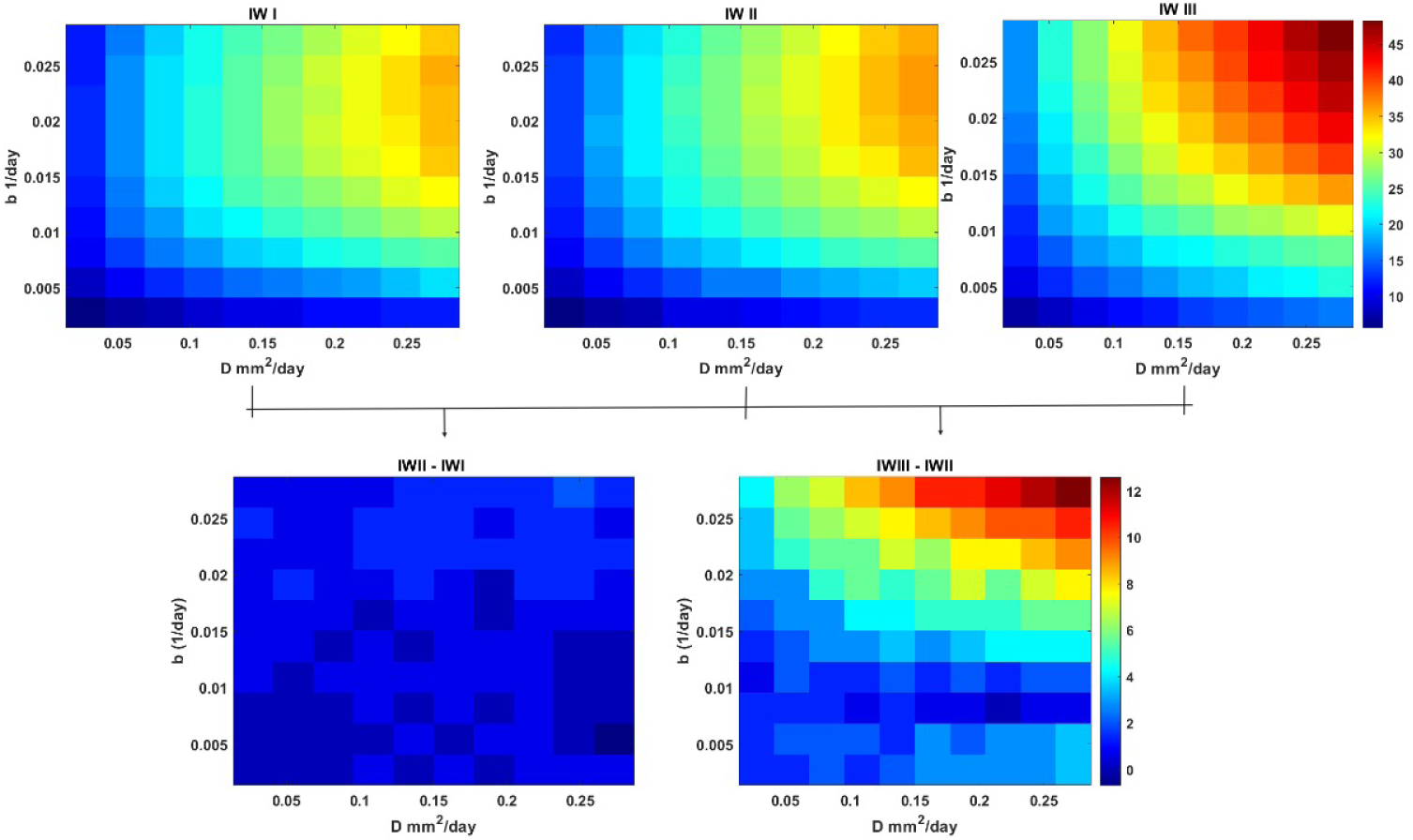
Simulation maps based on intrinsic infiltration and proliferation rates (*D ∈* [2.73*×*10*^−^*^3^, 2.73*×*10*^−^*^1^) and *b ∈* [2.73*×*10*^−^*^4^, 2.73*×*10*^−^*^2^) A) IW for the maps for different values of glioma oxygen consumption rates (*h*2 = [5.73 *×* 10*^−^*^4^, 5.73 *×* 10*^−^*^3^, 5.73 *×* 10*^−^*^2^] and B) is the differences between IWI-II and IW III-II.

Fig. 8 illustrated that under low rates of oxygen consumption, gliomas exhibited a tendency to grow and proliferate more than infiltrate. Remarkably, this pattern remained consistent even at moderate oxygen consumption rates. However, as the TME became increasingly hypoxic due to elevated oxygen consumption rates by glioma cells, there was a notable decrease in TS attributed to a shift towards invasiveness and reduced proliferation. This shift became particularly evident when transitioning from TSI to TSIII. Quantitatively, the percentage decrease in TS was recorded as -0.42% from the maximum value of TSI to TSII, followed by a more substantial decline of -3.5% from the maximum value of TSII to TSIII.

**Fig. 8.**
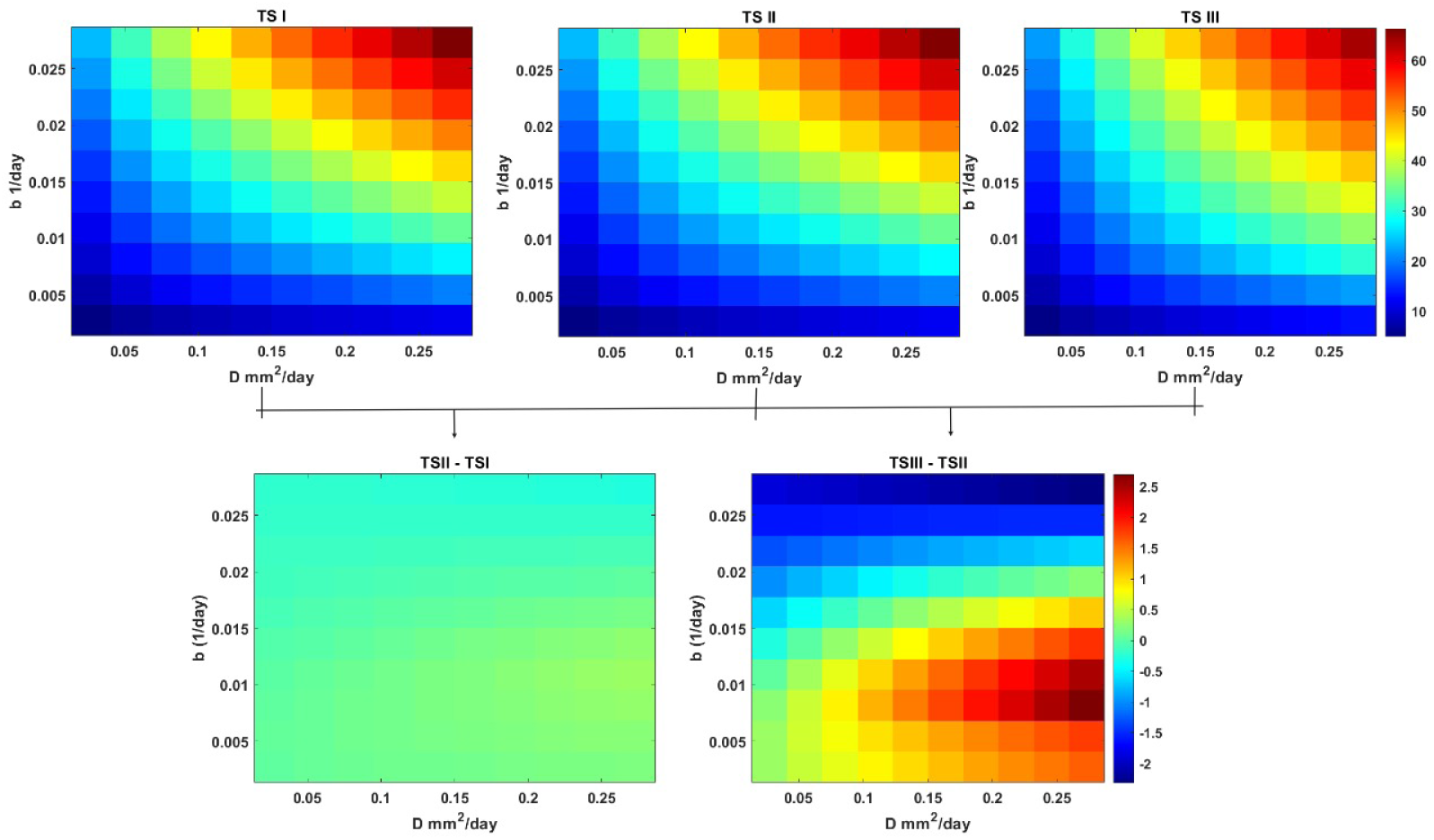
Simulation maps based on intrinsic infiltration and proliferation rates (*D ∈* [2.73*×*10*^−^*^3^, 2.73*×*10*^−^*^1^) and *b ∈* [2.73*×*10*^−^*^4^, 2.73*×*10*^−^*^2^) A) TS for the maps for different values of glioma oxygen consumption rates (*h*2 = [5.73 *×* 10*^−^*^4^, 5.73 *×* 10*^−^*^3^, 5.73 *×* 10*^−^*^2^] and B) is the differences between TS (I-II) and TS (III-II).

#### 3.1.2 Time dependant sensitivity analysis reveals the most effective parameter through the time to detection

Our time-dependent sensitivity analysis unveiled the pivotal role of the diffusion rate of glioma cells *D* as the most influential parameter affecting both quantities of interest, namely IW and TS. Notably, the diffusion rate demonstrated a robust positive correlation with TS and IW outputs. This correlation arose from the capacity of glioma cells to extend outward into the tumor core, where they can support proliferation, thus fostering tumor growth and invasion. Similarly, the tumor proliferation rate *b* emerged as another dominant positive factor influencing both IW and TS outputs—this observation aligned with established understanding, highlighting the significance of these parameters in shaping tumor behavior. The glioma oxygen consumption rate parameter (*h*_2_) emerged as a critical determinant, exhibiting a positive correlation with both IW and TS. Variations in glioma consumption rate could modulate tumor invasiveness at higher values and promote proliferation at lower values, delineating normoxic and hypoxic regions over time. In addition to these key parameters, the phenotypic regulation term (*t_n_*) exerted a notable positive effect on IW, particularly in later times, underscoring its role during the emergence of hypoxic tumor regions. Fig. 9 visually presented the time-dependent sensitivity analysis for the 11 parameters of the model, offering a comprehensive insight into their dynamic impact on tumor behavior over time.

**Fig. 9.**
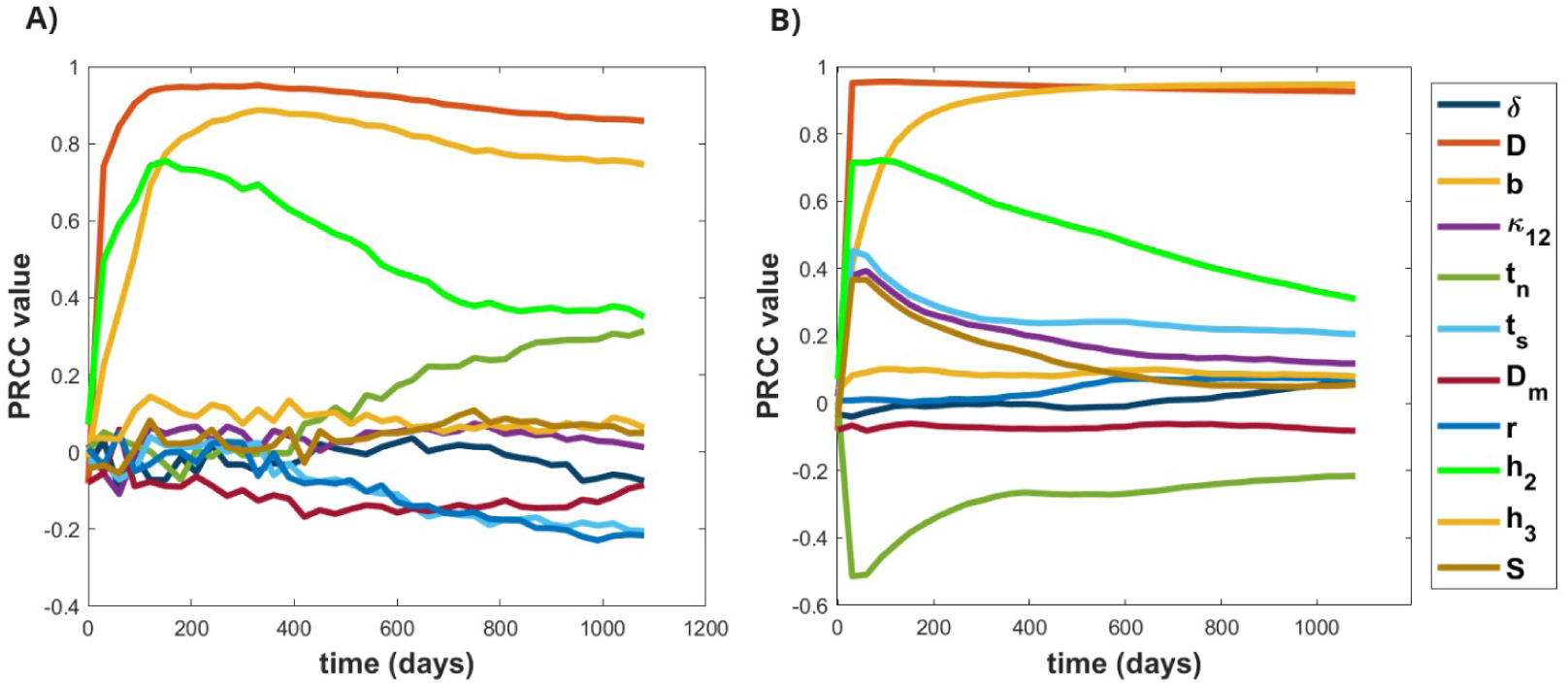
A) shows the PRCC sensitivity analysis for the IW and B) for the TS depicting the time-dependent analysis of 11 parameters and 500 samples to see their effects on the output.

### 3.2 The prognostic power of standard-of-care diagnostics and biopsy locus in predicting glioma recurrence

To address the question concerning the predictive potential of features within our study, we generated a synthetic dataset comprising 20,000 patient profiles. Primarily, this dataset underwent meticulous preprocessing, which laid the groundwork for an exhaustive analysis aimed at calculating correlation coefficients and mutual information metrics for each feature. The latter quantified even non-linear relationships among the variables. Following this, we partitioned the dataset into two primary categories: nodular and infiltrative gliomas. This stratification enabled us to assess the significance of specific features, such as macrophage ratios, Ki67 marker expressions, and tumor volumes as discerned in T1-weighted and FLAIR imaging modalities. Through this comprehensive investigation, our objective was to determine the impact of these variables on the overall predictive accuracy of the dataset, evaluated through mean squared error (MSE) and R-squared (*R*^2^) metrics.

#### 3.2.1 Localized macrophages at the edge: superior predictors in Glioma Analysis based on Preprocessing the data

Our preliminary analysis of the raw complete dataset revealed insightful correlations between features and the target variable. In Fig. 10A, the correlation coefficient heatmap illustrated the relationships among all features and the targets. Notably, tumor volumes at the time of detection demonstrated a strong correlation coefficient with the target, particularly with IW after 12 months post-resection. This positive correlation suggested that higher tumor volume values served as a robust predictor for IW after resection, peaking at the 12-month mark. In contrast, Fig. 9B, representing the mutual information plot, underscored a significant relationship between these features, hinting at the potential significance of initial tumor size as a predictor of its subsequent behavior and size upon recurrence.

**Fig. 10.**
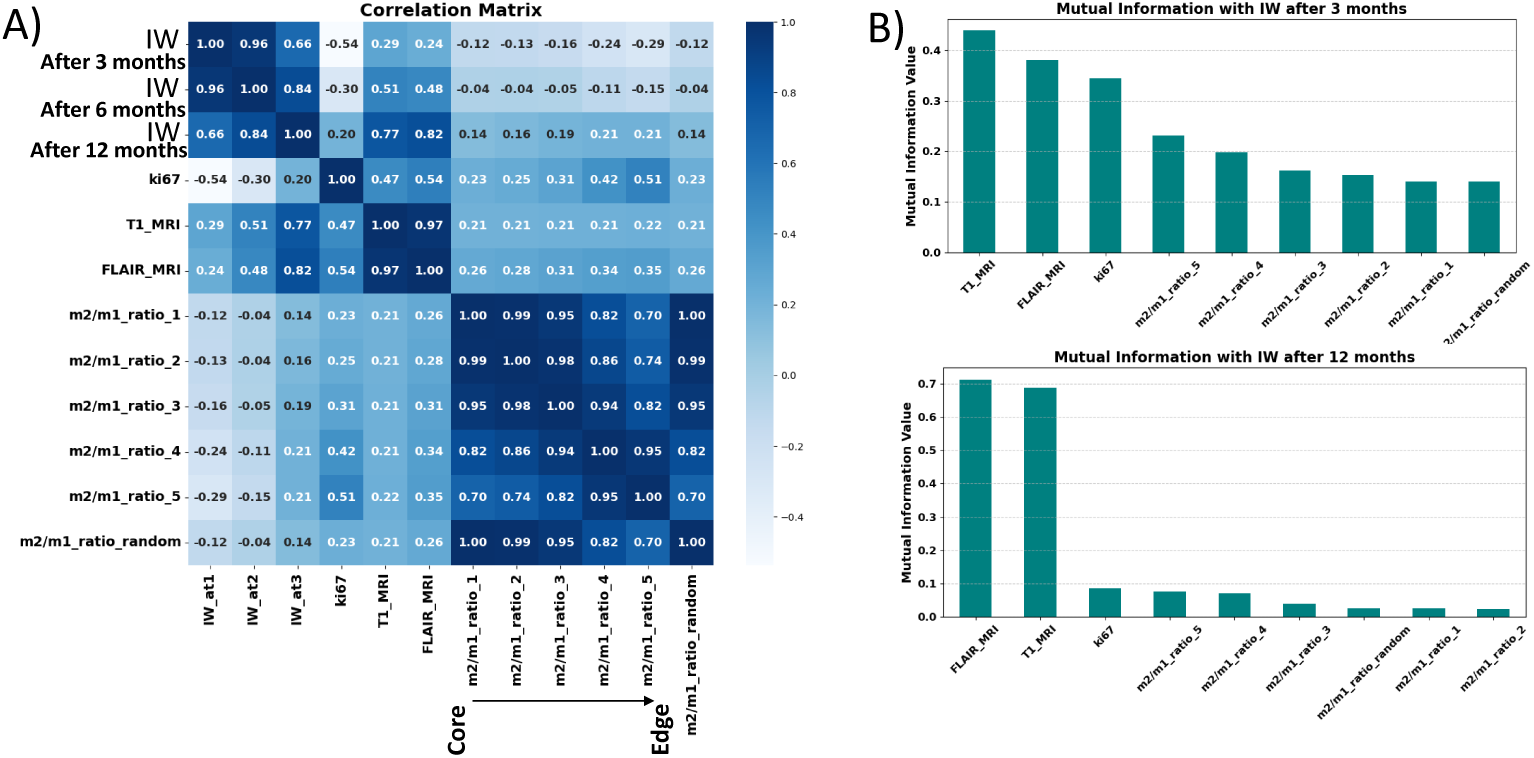
A) Correlation matrix heatmap for the raw dataset and the IW after 3, 6, and 12 months post resection (*IW*_a_*t* 1 depicts the IW after 3 months post resection and *IW*_a_*t*3 shows the IW after 12 months) B) Mutual information bar plot for the features and IW (3 and 12 months) T1 and FLAIR MRI volumes have the highest correlation and mutual information values to the target.

However, the behavior of the Ki67 marker presented an intriguing dynamic: negative values were observed for IW after 3 and 6 months, while a positive value emerged for IW after 12 months. This discrepancy suggested that in the early stages of tumor recurrence, a high Ki67 index might indicate rapid cell division, temporarily suppressing the tumor’s invasive characteristics as it focused on rebuilding mass rather than invading new territories. As the tumor matured, its relationship with the microenvironment evolved, potentially fostering a more conducive environment for infiltration. Similarly, due to the highly non-linear behavior of the system, macrophages exhibited a similar pattern as Ki67. They had a very low negative effect on IW after 3 and 6 months and a positive value for IW after 12 months post-resection. This behavior could be attributed to the shift in macrophage phenotype, which was crucial for understanding their changing impact on tumor infiltrative behavior.

Additionally, intriguing results highlighted that macrophages close to the tumor resection threshold (Edge region) exhibited higher values compared to macrophage ratios close to the tumor core. This pattern was particularly evident for IW after 12 months post-resection, suggesting that macrophages predominantly harbored a pro-tumor phenotype and might be more beneficial at the edge to facilitate increased invasiveness. Fig. 10 A, B presented the correlation heatmap and mutual information for the raw dataset, providing a comprehensive visual representation of the complex interplay between features and tumor behavior.

#### 3.2.2 Impact of noise on infiltration width predictive models for nodular and infiltrative tumors

In this section, we evaluate the effects of noise on predictive modeling for nodular and infiltrative tumors at 3 and 12 months post-resection. Fig. 11 presents the performance metrics, including MSE and *R*^2^, under various scenarios, encompassing different feature sets.

**Fig. 11.**
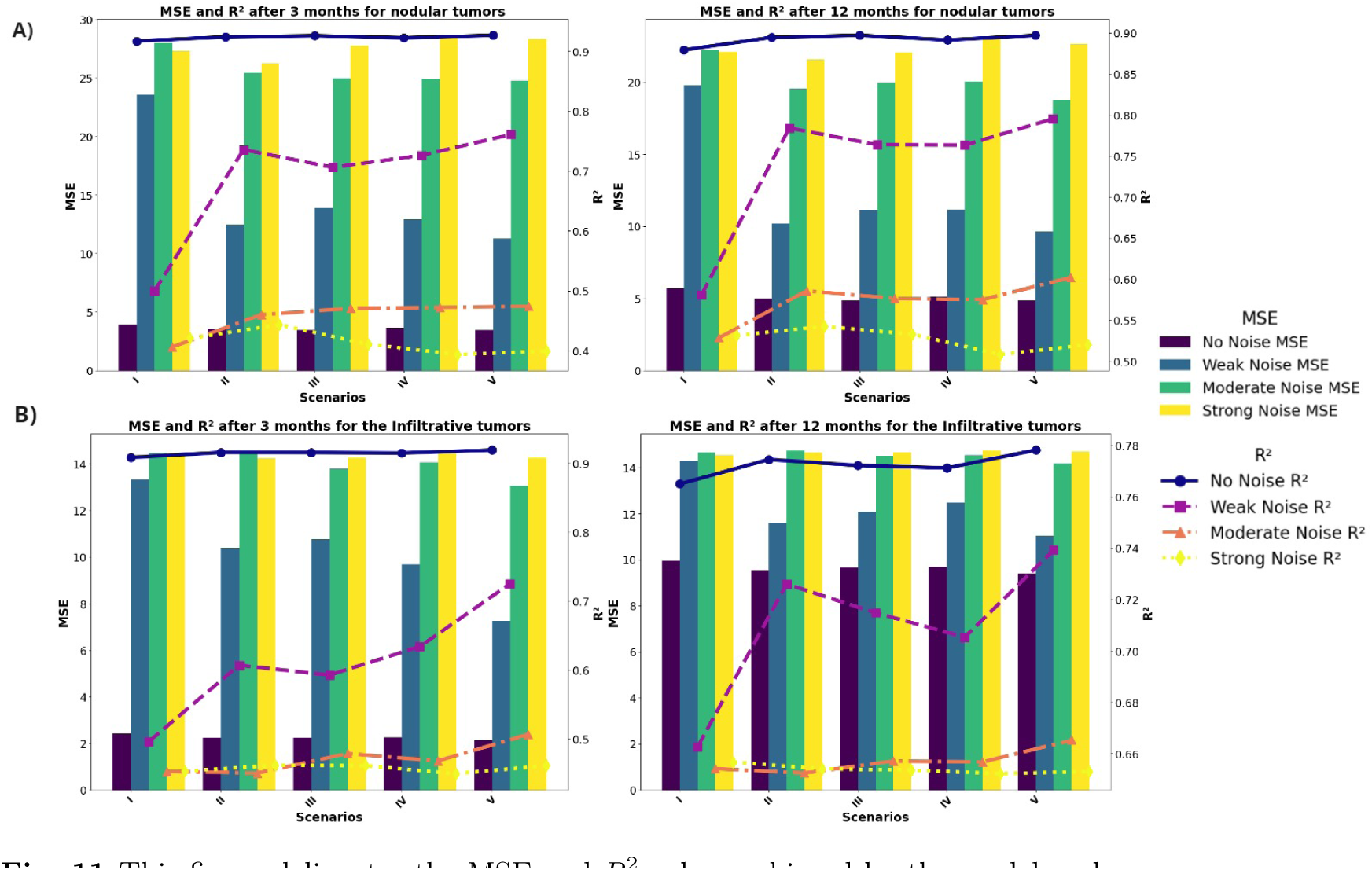
This figure delineates the MSE and *R*^2^ values achieved by the model under varying levels of data noise (No Noise, Weak Noise, Moderate Noise, Strong Noise) and across five distinct feature scenarios (I-V). Each scenario represents a combination of biological and imaging markers used to predict the output for 3 and 12 months post-resection.

##### Analysis for nodular tumors

We begin by assessing the predictive performance of nodular tumors across different noise levels. Notably, zero-noise conditions consistently yield the best outcomes across all scenarios. For instance, the model achieves an MSE of 3.90 and an *R*^2^ of 0.917 for nodular tumors three months post-resection. However, with the introduction of weak noise, performance deteriorates substantially, indicating a high sensitivity to noise. Incorporating macrophage ratios enhances model performance under noise-free conditions, with Scenario V displaying the most robust performance, demonstrating resilience against weak and moderate noise levels. Moving to 12 months post-resection, the results show a higher MSE for the zero noise cases compared to the dataset at 3 months post-resection. However, under weak noise, Scenario V demonstrates the most robustness (MSE: 9.66, *R*^2^: 0.795), indicating the benefits of including both core and edge macrophage ratios.

##### Analysis for infiltrative tumors

In contrast, infiltrative tumors exhibit a higher sensitivity to noise, with increased MSE and decreased *R*^2^ values under weak noise conditions. However, models with comprehensive features (Scenario V) display greater resilience against noise, particularly under weak and moderate noise levels. Despite the introduction of noise, Scenario V consistently yields the most accurate predictions, underscoring the importance of a broad feature set in enhancing model stability.

The comprehensive analysis highlights the significance of data quality in predictive accuracy, with nodular tumors showing greater vulnerability to noise compared to infiltrative tumors. Additionally, including macrophage ratios enhances predictive performance, particularly in scenarios with noise-free conditions. For both kinds of tumors, the feature set consisting of Ki67, T1, and FLAIR MRI, and macrophage ratios, at the core and edge of tumors, (Scenario V) offers the most accurate predictions

#### 3.2.3 Robustness of feature importance in predicting infiltration width post-resection

In this section, we investigate the robustness of feature importance in predicting IW post-resection across different data quality levels. Fig. 12 illustrates the feature importance for nodular and infiltrative datasets at both 3 and 12 months post-resection, considering varying levels of noise.

**Fig. 12.**
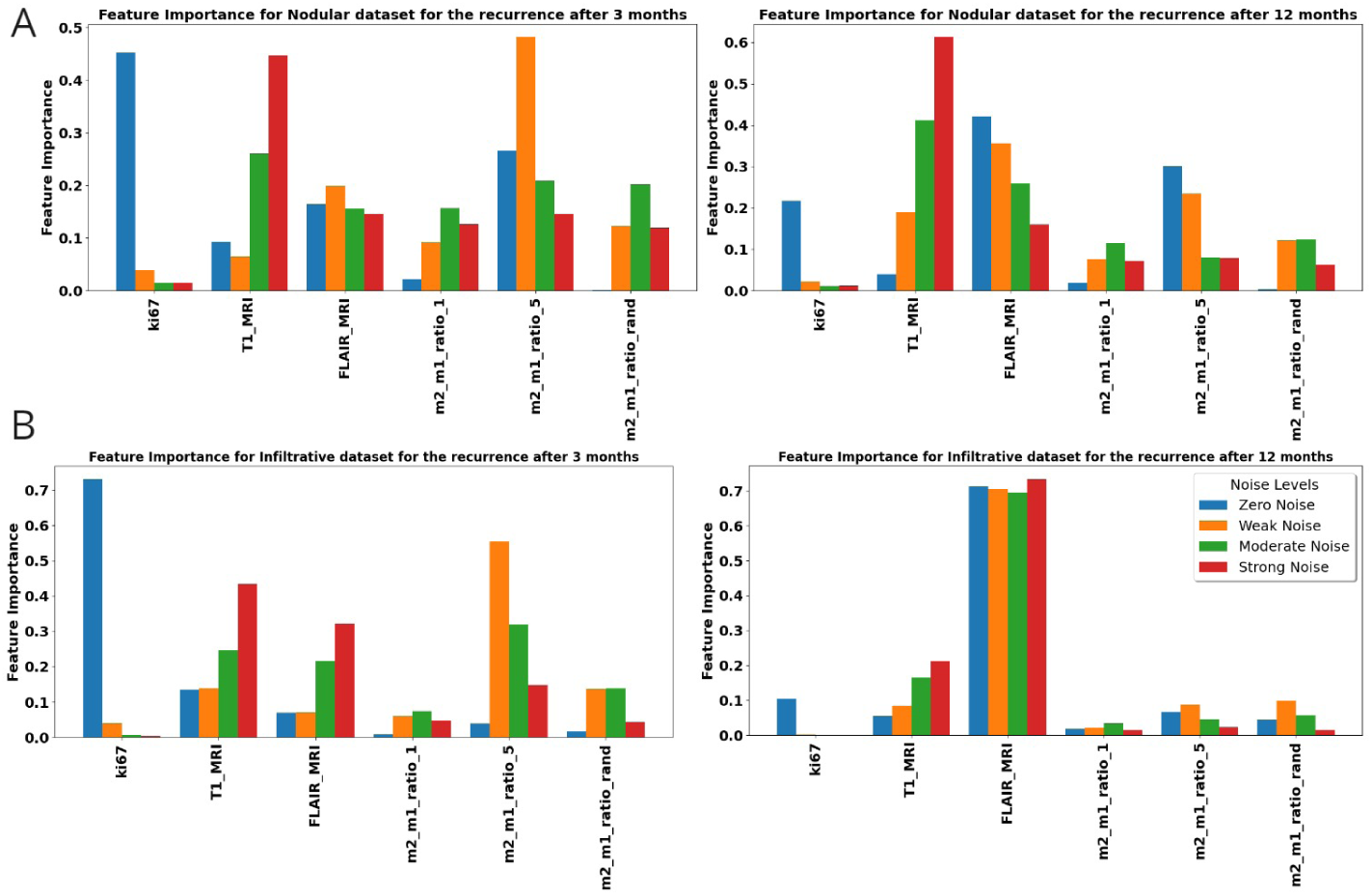
The bar chart shows feature importance scores for different features in a dataset, under four conditions: No Noise, Weak Noise, Moderate Noise, and Strong Noise for the nodular and infiltrative dataset 3, and 12 months post resection.

Initially, the Ki67 index emerges as a strong predictor in noise-free datasets, indicating its significance in tumor prognosis. However, the predictive reliability of Ki67 diminishes with the introduction of noise, highlighting its vulnerability to data perturbations. This observation underscores the heightened sensitivity of Ki67 to cellular proliferation rates, particularly in the early stages of tumor regrowth post-resection. This decline is likely attributed to the evolved state of the tumor at later stages, where the complexity of tumor biology extends beyond proliferation rates, rendering Ki67 a less definitive prognostic feature.

Furthermore, macrophage ratios, particularly the *m*2*/m*1 ratio at the tumor edge, exhibit stability across different noise levels, suggesting intrinsic characteristics of the tumor microenvironment’s response to resection. This stability underscores the potential utility of localized biopsies in predicting tumor behavior, even in scenarios with poor data quality. Overall, the analysis underscores the importance of robust feature selection and data quality management in enhancing the interpretability and reliability of predictive models for tumor prognosis.

## 4 Discussion

In this study, we developed a spatial mechanistic model that provided new insights into the dual phenotypic plasticity of glioma cells and macrophages, as well as their intricate interactions. Our model elucidated the nonlinear dynamics of these interactions and revealed how macrophages and oxygen availability influenced the plasticity of glioma cells through mechanisms governed by the ”go-or-grow” dichotomy. This dichotomy, which described the critical trade-off between tumor cell proliferation and migration, was captured effectively in our model, reflecting the complex biological behaviors of GBM as observed in clinical settings. The model highlighted how variations in oxygen consumption rate could lead to more infiltrative and less proliferative tumor phenotypes, aligning with previous studies that explored the impact of microenvironmental factors such as oxygen availability and mechanical stress on tumor dynamics [38, 37].

After exploring the parameter space via global sensitivity analysis, we used the model as a virtual reality to investigate the predictive power of available in silico biopsies and MRIs toward recurrences after resection. To thoroughly assess the predictive capabilities of the features generated from our model, we constructed a comprehensive synthetic dataset comprising 20,000 virtual patient profiles. This dataset underwent preprocessing to ensure data integrity and usability, incorporating variability through the application of Gaussian noise with three distinct standard deviations. We employed statistical techniques to derive correlation coefficients and mutual information metrics for each feature, then classified the dataset into nodular and infiltrative gliomas, representing two distinct patterns of brain tumor growth crucial for guiding clinical management and mathematical modeling.

Our results indicated that at the early stages of intratumoral vessel density (IW) post-resection (IW after 3 months), Ki67 emerged as the strongest predictive biomarker for the zero-noise dataset, reflecting the role of proliferative tumor cells at the early stage of relapse. However, even weak noise significantly diminished Ki67’s predictive power, highlighting the impact of technical and biological variability on biomarker assessment. Conversely, the *m*_2_*/m*_1_ ratio at the tumor edge exhibited high predictive power, especially for recurrence 3 months post-resection, suggesting the potential necessity of locus-specific biopsies in improving predictability. Interestingly, the latter was rather robust for different noise levels.

Our analysis also revealed that T1 and FLAIR MRI tumor volumes exhibited a stronger correlation with tumor recurrence at 12 months post-resection, indicating their enhanced predictive capacity for advanced recurrent tumors compared to early-stage ones. This could be explained since tumor scans set the initial conditions for any predictive model. In particular, MRI-based measurements could have been pivotal in stratifying patients based on their risk of late recurrence, thereby informing more tailored post-surgical interventions and monitoring strategies.

However, our study was limited by the inherent assumptions and simplifications required during the model development. The deterministic nature of our model might not have fully accounted for the stochastic variations seen in actual biological systems, despite introducing controlled noise to simulate this randomness. Additionally, we did not consider the impact of chemo- and radiotherapy on relapse, focusing instead on investigating the predictive power of generated features within a synthetic reality scenario.

In our study, we focused simplistically on the role of monocyte-derived macrophages, which circulated in the blood vessels and were recruited by tumor cells. The potential particularities of resident microglia were not included in our model.

Moreover, as hypoxia was acknowledged as a factor in our assumptions, we maintained a constant uniform value for the vasculature within the tumor microenvironment (TME) for simplification. We used synthetic classical MRI data, specifically FLAIR and T1Gd modalities, in our imaging approach. However, the MRI protocol could be expanded to include additional modalities such as PET, DTI, DWI, ADC, and SWI for each patient. These additional modalities could have provided a richer dataset and enhanced the information available for analyzing the patient cohort. In summary, our study offered valuable insights into the predictive power of available features in predicting tumor recurrence post-resection. Future studies, both in silico and experimental, could help validate and identify the predictive power of existing features, facilitating more personalized treatment strategies for glioma patients.

## Supporting information

https://tex.zih.tu-dresden.de/project/6602a9961ce17f4fcf9e3b27

## Acknowledgments

P.Sh. and H.H. have received funding from the Bundesministerium für Bildung und Forschung (BMBF) under grant agreement No. 031L0237C (MiEDGE project/ERACOSYSMED). H.H. would also like to acknowledge the support of the Volkswagen Stiftung for the “Life?” initiative (grant number 96732). Finally, HH acknowledges the support of the RIG-2023-051 grant from Khalifa University and the UAE-NIH Collaborative Research grant AJF-NIH-25-KU. We extend our thanks to Prof. Anna-Maria Pappa for her insightful help. E.W. received funding from the European Union, co-financed by tax revenues based on the budget adopted by the Saxon State Parliament, as part of an ESF Plus scholarship (grant number 100670474).

## Notes

### Competing Interest Statement

The authors have declared no competing interest.

## References

[1] Q. T. Ostrom, M. Price, C. Neff, G. Cioffi, K. A. Waite, C. Kruchko, J. S. Barnholtz-Sloan, Cbtrus statistical report: primary brain and other central nervous system tumors diagnosed in the united states in 2015–2019, Neuro-oncology 24 (Supplement 5) (2022) v1–v95.

[2] P. Wen, M. Weller, E. Lee, et al., Glioblastoma in adults: a sno and eano consensus review on current management and future directions, Neuro Oncol. (2020).

[3] J. J. Miller, L. N. Gonzalez Castro, S. McBrayer, M. Weller, T. Cloughesy, J. Portnow, O. Andronesi, J. S. Barnholtz-Sloan, B. G. Baumert, M. S. Berger, et al., Isocitrate dehydrogenase (idh) mutant gliomas: a society for neuro-oncology (sno) consensus review on diagnosis, management, and future directions, Neuro-oncology 25 (1) (2023) 4–25.

[4] R. Stupp, W. P. Mason, M. J. Van Den Bent, M. Weller, B. Fisher, M. J. Taphoorn, K. Belanger, A. A. Brandes, C. Marosi, U. Bogdahn, et al., Radiotherapy plus concomitant and adjuvant temozolomide for glioblastoma, New England journal of medicine 352 (10) (2005) 987–996.

[5] C. Fernandes, A. Costa, L. Osório, R. C. Lago, P. Linhares, B. Carvalho, C. Caeiro, Current standards of care in glioblastoma therapy, Exon Publications (2017) 197–241.

[6] H. A. Fine, New strategies in glioblastoma: exploiting the new biology, Clinical Cancer Research 21 (9) (2015) 1984–1988.

[7] D. Hanahan, Hallmarks of cancer: new dimensions, Cancer discovery 12 (1) (2022) 31–46.

[8] M. Nakada, D. Kita, T. Watanabe, Y. Hayashi, L. Teng, I. V. Pyko, J.-I. Hamada, Aberrant signaling pathways in glioma, Cancers 3 (3) (2011) 3242–3278.

[9] M. C. Tate, M. K. Aghi, Biology of angiogenesis and invasion in glioma, Neurotherapeutics 6 (3) (2009) 447–457.

[10] M. Onishi, T. Ichikawa, K. Kurozumi, I. Date, Angiogenesis and invasion in glioma, Brain tumor pathology 28 (2011) 13–24.

[11] A. A. Hardigan, J. D. Jackson, A. P. Patel, Surgical management and advances in the treatment of glioma, in: Seminars in Neurology, Thieme Medical Publishers, Inc., 2023.

[12] A. Claes, A. J. Idema, P. Wesseling, Diffuse glioma growth: a guerilla war, Acta neuropathologica 114 (2007) 443–458.

[13] A. Giese, R. Bjerkvig, M. Berens, M. Westphal, Cost of migration: invasion of malignant gliomas and implications for treatment, Journal of clinical oncology 21 (8) (2003) 1624–1636.

[14] J. Y. Kim, J. E. Park, Y. Jo, W. H. Shim, S. J. Nam, J. H. Kim, R.-E. Yoo, S. H. Choi, H. S. Kim, Incorporating diffusion-and perfusion-weighted mri into a radiomics model improves diagnostic performance for pseudoprogression in glioblastoma patients, Neuro-oncology 21 (3) (2019) 404–414.

[15] D. P. Barboriak, Z. Zhang, P. Desai, B. S. Snyder, Y. Safriel, R. C. McKinstry, F. Bokstein, G. Sorensen, M. R. Gilbert, J. L. Boxerman, Interreader variability of dynamic contrast-enhanced mri of recurrent glioblastoma: the multicenter acrin 6677/rtog 0625 study, Radiology 290 (2) (2019) 467–476.

[16] J. Wang, X. Yi, Y. Fu, P. Pang, H. Deng, H. Tang, Z. Han, H. Li, J. Nie, G. Gong, et al., Preoperative magnetic resonance imaging radiomics for predicting early recurrence of glioblastoma, Frontiers in Oncology 11 (2021) 769188.

[17] A. Giese, M. A. Loo, N. Tran, D. Haskett, S. W. Coons, M. E. Berens, Dichotomy of astrocytoma migration and proliferation, International journal of cancer 67 (2) (1996) 275–282.

[18] A. Giese, L. Kluwe, B. Laube, H. Meissner, M. E. Berens, M. Westphal, Migration of human glioma cells on myelin, Neurosurgery 38 (4) (1996) 755–764.

[19] H. Hatzikirou, D. Basanta, M. Simon, K. Schaller, A. Deutsch, ‘go or grow’: the key to the emergence of invasion in tumour progression?, Mathematical medicine and biology: a journal of the IMA 29 (1) (2012) 49–65.

[20] J. Godlewski, A. Bronisz, M. O. Nowicki, E. A. Chiocca, S. Lawler, microrna-451: A conditional switch controlling glioma cell proliferation and migration, Cell cycle 9 (14) (2010) 2814–2820.

[21] E. Höring, P. N. Harter, J. Seznec, J. Schittenhelm, H.-J. Bühring, S. Bhattacharyya, E. von Hattingen, C. Zachskorn, M. Mittelbronn, U. Naumann, The “go or grow” potential of gliomas is linked to the neuropeptide processing enzyme carboxypeptidase e zand mediated by metabolic stress, Acta neuropathologica 124 (2012) 83–97.

[22] S. D. Wang, P. Rath, B. Lal, J.-P. Richard, Y. Li, C. R. Goodwin, J. Laterra, S. Xia, Ephb2 receptor controls proliferation/migration dichotomy of glioblastoma by interacting with focal adhesion kinase, Oncogene 31 (50) (2012) 5132–5143.

[23] T. Stylianopoulos, L. L. Munn, R. K. Jain, Reengineering the physical microenvironment of tumors to improve drug delivery and efficacy: from mathematical modeling to bench to bedside, Trends in cancer 4 (4) (2018) 292–319.

[24] D. Hambardzumyan, D. H. Gutmann, H. Kettenmann, The role of microglia and macrophages in glioma maintenance and progression, Nature neuroscience 19 (1) (2016) 20–27.

[25] I. Bettinger, S. Thanos, W. Paulus, Microglia promote glioma migration, Acta neuropathologica 103 (2002) 351–355.

[26] D. F. Quail, J. A. Joyce, Microenvironmental regulation of tumor progression and metastasis, Nature medicine 19 (11) (2013) 1423–1437.

[27] A. R. Pombo Antunes, I. Scheyltjens, F. Lodi, J. Messiaen, A. Antoranz, J. Duerinck, D. Kancheva, L. Martens, K. De Vlaminck, H. Van Hove, et al., Single-cell profiling of myeloid cells in glioblastoma across species and disease stage reveals macrophage competition and specialization, Nature neuroscience 24 (4) (2021) 595–610.

[28] W. Yin, Y.-F. Ping, F. Li, S.-Q. Lv, X.-N. Zhang, X.-G. Li, Y. Guo, Q. Liu, T.-R. Li, L.-Q. Yang, et al., A map of the spatial distribution and tumour-associated macrophage states in glioblastoma and grade 4 idh-mutant astrocytoma, The Journal of pathology 258 (2) (2022) 121–135.

[29] P. Shojaee, F. Mornata, A. Deutsch, M. Locati, H. Hatzikirou, The impact of tumor associated macrophages on tumor biology under the lens of mathematical modelling: A review, Frontiers in Immunology 13 (2022) 1050067.

[30] A. Buonfiglioli, D. Hambardzumyan, Macrophages and microglia: the cerberus of glioblastoma, Acta neuropathologica communications 9 (1) (2021) 1–21.

[31] N. Ochocka, P. Segit, K. A. Walentynowicz, K. Wojnicki, S. Cyranowski, J. Swatler, J. Mieczkowski, B. Kaminska, Single-cell rna sequencing reveals functional heterogeneity of glioma-associated brain macrophages, Nature communications 12 (1) (2021) 1151.

[32] S. Gordon, Alternative activation of macrophages, Nature reviews immunology 3 (1) (2003) 23–35.

[33] Y. Han, X. Wang, K. Xia, T. Su, A novel defined hypoxia-related gene signature to predict the prognosis of oral squamous cell carcinoma, Annals of Translational Medicine 9 (20) (2021).

[34] M. Yang, D. McKay, J. W. Pollard, C. E. Lewis, Diverse functions of macrophages in different tumor microenvironments, Cancer research 78 (19) (2018) 5492–5503.

[35] A.-T. Henze, M. Mazzone, et al., The impact of hypoxia on tumor-associated macrophages, The Journal of clinical investigation 126 (10) (2016) 3672–3679.

[36] R. Bai, Y. Li, L. Jian, Y. Yang, L. Zhao, M. Wei, The hypoxia-driven crosstalk between tumor and tumor-associated macrophages: mechanisms and clinical treatment strategies, Molecular Cancer 21 (1) (2022) 177.

[37] P. Mascheroni, J. C. López Alfonso, M. Kalli, T. Stylianopoulos, M. Meyer-Hermann, H. Hatzikirou, On the impact of chemo-mechanically induced phenotypic transitions in gliomas, Cancers 11 (5) (2019) 716.

[38] J. Alfonso, A. Köhn-Luque, T. Stylianopoulos, F. Feuerhake, A. Deutsch, H. Hatzikirou, Why one-size-fits-all vaso-modulatory interventions fail to control glioma invasion: in silico insights, Scientific reports 6 (1) (2016) 1–15.

[39] G. E. Mahlbacher, K. C. Reihmer, H. B. Frieboes, Mathematical modeling of tumor-immune cell interactions, Journal of Theoretical Biology 469 (2019) 47–60.

[40] R. Eftimie, J. L. Bramson, D. J. Earn, Interactions between the immune system and cancer: a brief review of non-spatial mathematical models, Bulletin of mathematical biology 73 (2011) 2–32.

[41] J. A. Bull, H. M. Byrne, The hallmarks of mathematical oncology, Proceedings of the IEEE 110 (5) (2022) 523–540.

[42] M. Kuznetsov, J. Clairambault, V. Volpert, Improving cancer treatments via dynamical biophysical models, Physics of life reviews 39 (2021) 1–48.

[43] K. Böttger, H. Hatzikirou, A. Chauviere, A. Deutsch, Investigation of the migration/proliferation dichotomy and its impact on avascular glioma invasion, Mathematical Modelling of Natural Phenomena 7 (1) (2012) 105–135.

[44] G. Wang, K. Zhong, Z. Wang, Z. Zhang, X. Tang, A. Tong, L. Zhou, Tumor-associated microglia and macrophages in glioblastoma: From basic insights to therapeutic opportunities, Frontiers in Immunology 13 (2022) 964898.

[45] J. K. Andersen, H. Miletic, J. A. Hossain, Tumor-associated macrophages in gliomas—basic insights and treatment opportunities, Cancers 14 (5) (2022) 1319.

[46] J. Liu, X. Geng, J. Hou, G. Wu, New insights into m1/m2 macrophages: key modulators in cancer progression, Cancer Cell International 21 (1) (2021) 1–7.

[47] F. Zhang, N. Parayath, C. Ene, S. Stephan, A. Koehne, M. Coon, E. Holland, M. Stephan, Genetic programming of macrophages to perform antitumor functions using targeted mrna nanocarriers, Nature communications 10 (1) (2019) 3974.

[48] B. Muz, P. de la Puente, F. Azab, A. Kareem Azab, The role of hypoxia in cancer progression, angiogenesis, metastasis, and resistance to therapy, Hypoxia (2015) 83–92.

[49] P. Carmeliet, R. K. Jain, Angiogenesis in cancer and other diseases, nature 407 (6801) (2000) 249–257.

[50] S. Magri, B. Musca, L. Pinton, E. Orecchini, M. L. Belladonna, C. Orabona, C. Bonaudo, F. Volpin, P. Ciccarino, V. Baro, et al., The immunosuppression pathway of tumor-associated macrophages is controlled by heme oxygenase-1 in glioblastoma patients, International Journal of Cancer 151 (12) (2022) 2265–2277.

[51] H. J. Knowles, A. L. Harris, Macrophages and the hypoxic tumour microenvironment, Front Biosci 12 (5) (2007) 4298–4314.

[52] M. Badoual, C. Gerin, C. Deroulers, B. Grammaticos, J.-F. Llitjos, C. Oppenheim, P. Varlet, J. Pallud, Oedema-based model for diffuse low-grade gliomas: application to clinical cases under radiotherapy, Cell proliferation 47 (4) (2014) 369–380.

[53] K. R. Swanson, R. C. Rockne, J. Claridge, M. A. Chaplain, E. C. Alvord Jr, A. R. Anderson, Quantifying the role of angiogenesis in malignant progression of gliomas: in silico modeling integrates imaging and histology, Cancer research 71 (24) (2011) 7366–7375.

[54] H. L. Harpold, E. C. Alvord Jr, K. R. Swanson, The evolution of mathematical modeling of glioma proliferation and invasion, Journal of Neuropathology & Experimental Neurology 66 (1) (2007) 1–9.

[55] S. E. Eikenberry, T. Sankar, M. C. Preul, E. J. Kostelich, C. Thalhauser, Y. Kuang, Virtual glioblastoma: growth, migration and treatment in a three-dimensional mathematical model, Cell proliferation 42 (4) (2009) 511–528.

[56] J. McDaniel, E. Kostelich, Y. Kuang, J. Nagy, M. C. Preul, N. Z. Moore, N. L. Matirosyan, Data assimilation in brain tumor models, Mathematical methods and models in biomedicine (2013) 233–262.

[57] R. Eftimie, C. Barelle, Mathematical investigation of innate immune responses to lung cancer: The role of macrophages with mixed phenotypes, Journal of Theoretical Biology 524 (2021) 110739.

[58] W. E. Hoffman, F. T. Charbel, G. Edelman, K. Hannigan, J. I. Ausman, Brain tissue oxygen pressure, carbon dioxide pressure and ph during ischemia, Neurological research 18 (1) (1996) 54–56.

[59] A. Carreau, B. E. Hafny-Rahbi, A. Matejuk, C. Grillon, C. Kieda, Why is the partial oxygen pressure of human tissues a crucial parameter? small molecules and hypoxia, Journal of cellular and molecular medicine 15 (6) (2011) 1239–1253.

[60] I. Stamper, M. Owen, P. Maini, H. Byrne, Oscillatory dynamics in a model of vascular tumour growth-implications for chemotherapy, Biology direct 5 (1) (2010) 1–17.

[61] A. Matzavinos, C.-Y. Kao, J. E. F. Green, A. Sutradhar, M. Miller, A. Friedman, Modeling oxygen transport in surgical tissue transfer, Proceedings of the National Academy of Sciences 106 (29) (2009) 12091–12096.

[62] G. Powathil, M. Kohandel, M. Milosevic, S. Sivaloganathan, et al., Modeling the spatial distribution of chronic tumor hypoxia: implications for experimental and clinical studies, Computational and mathematical methods in medicine 2012 (2012).

[63] C. D. Eggleton, T. K. Roy, A. S. Popel, Predictions of capillary oxygen transport in the presence of fluorocarbon additives, American Journal of Physiology-Heart and Circulatory Physiology 275 (6) (1998) H2250–H2257.

[64] D. Goldman, A. S. Popel, A computational study of the effect of capillary network anastomoses and tortuosity on oxygen transport, Journal of theoretical biology 206 (2) (2000) 181–194.

[65] C. J. Kelly, M. Brady, A model to simulate tumour oxygenation and dynamic [18f]-fmiso pet data, Physics in Medicine & Biology 51 (22) (2006) 5859.

[66] D. R. Grimes, C. Kelly, K. Bloch, M. Partridge, A method for estimating the oxygen consumption rate in multicellular tumour spheroids, Journal of The Royal Society Interface 11 (92) (2014) 20131124.

[67] P. Vaupel, F. Kallinowski, P. Okunieff, Blood flow, oxygen and nutrient supply, and metabolic microenvironment of human tumors: a review, Cancer research 49 (23) (1989) 6449–6465.

[68] Z. Neufeld, W. von Witt, D. Lakatos, J. Wang, B. Hegedus, A. Czirok, The role of allee effect in modelling post resection recurrence of glioblastoma, PLoS computational biology 13 (11) (2017) e1005818.

[69] J.-S. Guo, Y. Morita, Entire solutions of reaction-diffusion equations and an application to discrete diffusive equations, Discrete and Continuous Dynamical Systems 12 (2) (2004) 193–212.

[70] K. Swanson, E. Alvord Jr, J. Murray, Virtual resection of gliomas: effect of extent of resection on recurrence, Mathematical and computer modelling 37 (11) (2003) 1177–1190.

[71] S. Marino, I. B. Hogue, C. J. Ray, D. E. Kirschner, A methodology for performing global uncertainty and sensitivity analysis in systems biology, Journal of theoretical biology 254 (1) (2008) 178–196.

[72] P. Macklin, M. E. Edgerton, A. M. Thompson, V. Cristini, Patient-calibrated agent-based modelling of ductal carcinoma in situ (dcis): from microscopic measurements to macroscopic predictions of clinical progression, Journal of theoretical biology 301 (2012) 122–140.

[73] X. Gao, J. T. McDonald, L. Hlatky, H. Enderling, Acute and fractionated irradiation differentially modulate glioma stem cell division kinetics, Cancer research 73 (5) (2013) 1481–1490.

