## Supplementary material for "The Role of Biopsy Position and Tumor-Associated Macrophages for Predictions on Recurrence of Malignant Gliomas: An In Silico Study": https://tex.zih.tu-dresden.de/project/6602a9961ce17f4fcf9e3b27: supp Material.pdf

### 1 Supplementary Material

#### 1.1 Stability Analysis of Glioma-Free Steady States

In the following, we conduct a stability analysis of the model that is proposed in the main text which is parameterized by the equations

$$
\begin{aligned}
\frac{\partial \rho_m}{\partial t} &= D\nabla^2 \rho_m - f_{mp}\rho_m + f_{pm}\rho_p - \delta m_1 \rho_m, \\
\frac{\partial \rho_p}{\partial t} &= b\rho_p(1-(\rho_m+\rho_p)/K) + f_{mp}\rho_m - f_{pm}\rho_p - \delta m_1 \rho_p, \\
\frac{\partial m_1}{\partial t} &= D_m\nabla^2 m_1 + S\frac{\rho_m+\rho_p}{\rho_m+\rho_p+K_p} - \kappa_{1,2} m_1(\rho_m+\rho_p) + \kappa_{2,1} m_2 - d_1 m_1, \\
\frac{\partial m_2}{\partial t} &= D_m\nabla^2 m_2 + \kappa_{1,2} m_1(\rho_m+\rho_p) - \kappa_{2,1} m_2 \\
&\quad + rH(n-n_{cr}) m_2 (1-\frac{(m_1+m_2)}{K_m}) - d_2 m_2, \\
\frac{\partial n}{\partial t} &= D_n\nabla^2 n + h_1 v(n_0-n) - h_2(\rho_m+\rho_p)n - h_3(m_1+m_2)n,
\end{aligned}
$$

[*] Center for Interdisciplinary Digital Sciences (CIDS), Department Information Services and High-Performance Computing (ZIH), Dresden University of Technology, 01062 Dresden, Germany

[†] Department of Neuropathology, Institute for Pathology, Hannover Medical School, Hannover, Germany

[‡] Institute for Neuropathology, University Clinic Freiburg, Freiburg, Germany

[§] Mathematics Department, Khalifa University, Abu Dhabi, UAE.

where

$$f_{pm} = t_s(\frac{m_2}{K}) + t_n(\gamma_n - n),$$
$$f_{mp} = t_n n$$

and

$$H(n - n_{cr}) = 1 - \frac{1}{1 + e^{-2\theta(n-n_{cr})}}.$$

To be more precise we perform a linear stability analysis of the diffusion-free (ODE) system ($D = D_m = D_n = 0$) following the terminology of [1]. The steady states are given by the solutions to the system of equations

$$\frac{\partial \rho_m}{\partial t} = \frac{\partial \rho_p}{\partial t} = \frac{\partial m_1}{\partial t} = \frac{\partial m_2}{\partial t} = \frac{\partial n}{\partial t} \stackrel{!}{=} 0. \tag{1}$$

We will only consider steady states as the solutions to (1) for which $\rho_m, \rho_p, m_1, m_2, n$ since only these have a biological meaning. For the 5D nonlinear system we are investigating, deriving analytic solutions for (1) is challenging. Additionally, determining the stability at the steady states requires determining the sign of the eigenvalues of the Jacobian matrix, which are given by the roots of a polynomial of degree 5. In general, there is no explicit formula for the roots of such polynomials. To reduce the complexity and to still ensure a biomedical meaning of the analysis we investigate the glioma-free steady states ($\rho_m = \rho_p = 0$) of the system. The results will help us to understand under which conditions the tumor will eventually die out over time.

The complexity of finding roots is further amplified by the continuous version of the Heaviside function $H$ which depends on the exponential function. Due to this, we will later separately analyze the system in the hypoxic ($H \approx 1$) and normoxic ($H \approx 0$) region.

We use the parameter values or ranges respectively as they are given in the main text (cf. Table 1).

Hence, throughout the stability analysis we will keep the following parameters fixed:

$$K, K_m, K_p, S, \delta, d_1, d_2, n_0, K_p, v, h_1, \theta.$$

On the other side we have ranges for the values of

$$D, b, \delta, t_s, t_n, r, h_2, h_3, \kappa_{1,2}, \kappa_{2,1}.$$

For convenience we will suppress the dependency of functions of all these parameters in the notation and only highlight the dependency on some of them if needed. To further reduce complexity we will occasionally assume that $t_s + t_n = 1$. Some of the data in the main text is generated for $t_s = t_n = 0.5$ which is just a special case of this assumption.

**Table 1** Parameter values and specific ranges in the model simulations

| Parameter | Value | Unit | Description (source) |
|---|---|---|---|
| $D$ | $[2.73\times10^{-3}, 2.73\times10^{-1}]$ | $mm^2day^{-1}$ | Diffusion rate of glioma cells [2, 3, 4] |
| $b$ | $[2.73\times10^{-4}, 2.73\times10^{-2}]$ | $day^{-1}$ | Glioma cell proliferation rate [2, 3, 4] |
| $\delta$ | $2\times10^{-4}$ | $day(cells)^{-1}$ | Glioma cell death rate induced by macrophages |
| $K$ | $10^2$ | $cells(mm)^{-1}$ | Tumor carrying capacity [5, 6] |
| $K_m$ | $10^2\times25\%$ | $cells(mm)^{-1}$ | macrophage carrying capacity [7] |
| $t_s$ | $[0, 0.5]$ | - | $m_1$ transition rate (Estimated) |
| $t_n$ | $[0, 0.5]$ | - | nutrient transition rate (Estimated) |
| $d_1$ | 0.87 | $day^{-1}$ | $m_1$ death rate [8] |
| $d_2$ | 0.1 | $day^{-1}$ | $m_2$ death rate [8] |
| $r$ | $[0.487, 0.88]$ | $day^{-1}$ | Macrophage division rate [8] |
| $n_0$ | 1 | $nmol(day)^{-1}$ | Physiological oxygen concentration [9, 10] |
| $n_{cr}$ | 0.25 | $nmol(day)^{-1}$ | Critical oxygen concentration |
| $K_p$ | $0.5\times K$ | - | saturation rate (Estimated) |
| $D_m$ | $[1\times10^{-3}, 1\times10^{-1}]$ | $mm^2day^{-1}$ | Macrophage diffusion rate (Estimated) |
| $D_n$ | $1.51\times10^2$ | $mm^2day^{-1}$ | Oxygen diffusion rate [11, 12, 13] |
| $h1$ | $3.37\times10^{-1}$ | $day^{-1}$ | Oxygen supply rate [14, 15, 16] |
| $h2$ | $[5.73\times10^{-3}, 1.14\times10^{-1}]$ | $mm(cell\cdot day)^{-1}$ | Glioma cell oxygen consumption rate [17, 18] |
| $h3$ | $[1\times10^{-3}, 1\times10^{-1}]$ | $mm(cell\cdot day)^{-1}$ | Oxygen consumption rate by macrophages (Estimated) |
| $v$ | 1 | $cells(mm)^{-1}$ | Vasculature coefficient |
| $\kappa_{1,2}$ | $[10^{-5}, 10^{-2}]$ | $(day\cdot cells)^{-1}$ | M1 to M2 switch rate [8] |
| $\kappa_{2,1}$ | $[10^{-10}, 10^{-9}]$ | $(day\cdot cells)^{-1}$ | M2 to M1 switch rate [8] |
| $S$ | $[0.1, 0.5]\times K_m$ | $cells(mm)^{-1}$ | Recruitment rate (estimated) |
| $\theta$ | 10 | $-$ | Slope parameter of the continuous Heaviside function $H$ |
| $\gamma_n$ | 1.01 | $-$ | Regularization term |

For the sake of clarity, we will mostly omit substitution of the fixed variables by their respective values. For the computations throughout we used [19].

For later reference, we write

$$F(\rho_m,\rho_p,m_1,m_2,n) := \begin{bmatrix} -f_{mp}\rho_m + f_{pm}\rho_p - \delta m_1\rho_m \\ b\rho_p(1-(\rho_m+\rho_p)/K) + f_{mp}\rho_m - f_{pm}\rho_p - \delta m_1\rho_p \\ S\frac{\rho_m+\rho_p}{\rho_m+\rho_p+K_p} - \kappa_{1,2}m_1(\rho_m+\rho_p) + \kappa_{2,1}m_2 - d_1m_1 \\ \kappa_{1,2}m_1(\rho_m+\rho_p) - \kappa_{2,1}m_2 + rH(n-n_{cr})m_2(1-\frac{(m_1+m_2)}{K_m}) - d_2m_2 \\ h_1v(n_0-n) - h_2(\rho_m+\rho_p)n - h_3(m_1+m_2)n. \end{bmatrix}$$

##### 1.1.1 Reminder of Properties of Parabolas

Parabolas will play an important role in our investigation of the eigenvalues of the jacobian of the system at the steady states.

A parabola

$$f(x) = ax^2 + bx + c, x \in \mathbb{R}, b, c \in \mathbb{R}, a \in \mathbb{R}\setminus\{0\}$$

has vertex

$$P_{ext} = (x_{ext}, f(x_{ext})) = \left(-\frac{b}{2a}, c - \frac{b^2}{4a}\right)$$

that is a minimum point of $f$ if $a > 0$ and a maximum point of $f$ if $a < 0$.

We also have

$$\begin{cases} a < 0 \implies & \begin{cases} x < x_{est} \implies f \text{ monotonously increasing} \\ x > x_{est} \implies f \text{ monotonously decreasing} \end{cases} \\ a > 0 \implies & \begin{cases} x < x_{est} \implies f \text{ monotonously decreasing} \\ x > x_{est} \implies f \text{ monotonously increasing} \end{cases} \end{cases} \tag{2}$$

which we will later exploit for a lot of estimates.

##### 1.1.2 Trivial Steady State

To begin the analysis we start with the trivial steady state of the system that is given by

$$x^*_{triv} = [0, 0, 0, 0, n_0].$$

This is the only steady state of the macrophage-free system ($m1 = m2 = 0$) for our parameter values.

The Jacobian of the system at $x^*_{triv}$ is given by

$$\nabla F(x^*_{triv}) =$$
$$\begin{bmatrix} -n_0 \cdot t_n & (\gamma_n - n_0) \cdot t_n & 0 & 0 & 0 \\ n_0 \cdot t_n & -(\gamma_n - n_0) \cdot t_n + b & 0 & 0 & 0 \\ S/K_p & S/K_p & -d_1 & \kappa_{2,1} & 0 \\ 0 & 0 & 0 & H(n_0 - n_{cr})r - d_2 - \kappa_{2,1} & 0 \\ -h_2 \cdot n_0 & -h_2 \cdot n_0 & -h_3 \cdot n_0 & -h_3 \cdot n_0 & -h_1 \cdot v \end{bmatrix}$$

and the eigenvalues are

$$\begin{bmatrix} \lambda_1 \\ \lambda_2 \\ \lambda_3 \\ \lambda_4 \\ \lambda_5 \end{bmatrix} := \begin{bmatrix} -1/2 \cdot \gamma_n \cdot t_n + 1/2 \cdot b - 1/2 \cdot \sqrt{f(t_n, b)} \\ -1/2 \cdot \gamma_n \cdot t_n + 1/2 \cdot b + 1/2 \cdot \sqrt{f(t_n, b)} \\ -h_1 \cdot v \\ H(n_0 - n_{cr}) \cdot r - d_2 - \kappa_{2,1} \\ -d_1 \end{bmatrix}.$$

where

$$f(t_n, b) := \gamma_n^2 \cdot t_n^2 + b^2 - 2 \cdot b \cdot (\gamma_n - 2 \cdot n_0) \cdot t_n$$

**Signs of** $\mathrm{Re}(\lambda_3), \mathrm{Re}(\lambda_5)$

Since the parameters $h_1, v$ and $d_1$ are positive, we have

$$\mathrm{Re}(\lambda_3), \mathrm{Re}(\lambda_5) < 0.$$

**Sign of** $\mathrm{Re}(\lambda_4)$

We have $H(n_0 - n_{cr}) \approx 3.059 \cdot 10^{-7}$, so

$$\mathrm{Re}(\lambda_4) \leq H(n_0 - n_{cr}) \cdot r - d_2 - \kappa_{2,1} \leq 3.06 \cdot 10^{-7} - d_2 - \kappa_{2,1} < 0.$$

**Signs of** $\mathrm{Re}(\lambda_1), \mathrm{Re}(\lambda_2)$

For investigating $\lambda_1$ and $\lambda_2$ we use the results collected in 1.1.1.

The function $f(t_n, b)$ describes a parabola in $b$ with vertex

$$\left((\gamma_n - 2 \cdot n_0) \cdot t_n, \gamma_n^2 \cdot t_n^2 - (\gamma_n - 2 \cdot n_0)^2 \cdot t_n^2\right).$$

We first want to show that $\sqrt{f}$ is real, so that $\lambda_1$ and $\lambda_2$ are real numbers.

For this we need that

$$\gamma_n^2 \cdot t_n^2 \geq (\gamma_n - 2 \cdot n_0)^2 \cdot t_n^2$$

which reduces to the constraint

$$\|\gamma_n\| \geq \|\gamma_n - 2 \cdot n_0\|.$$

This is satisfied for our specific values of $\gamma_n = 1.01$ and $n_0 = 1$, hence $f \geq 0$ and $\sqrt{f}$ are real.

To find some bounds for $\lambda_1$ and $\lambda_2$ we observe that the function $f(t_n, b)$ also defines a parabola in $t_n$ that attains its global minimum at

$$t_n = \tfrac{b \cdot (\gamma_n - 2 \cdot n_0)}{\gamma_n t_n} < 0.$$

Since the minima of the parabola in $b$ and in $t_n$ are both less than zero, the square root $\sqrt{f(t_n, b)}$ is monotonously increasing in $b$ and in $t_n$ for $b \geq 0$ and $t_n \geq 0$. In particular $\sqrt{f(t_n, b)}$ is monotonously increasing in $b$ and in $t_n$ for our possible $b$ and $t_n$ values.

Hence we get

$$\begin{aligned}\lambda_1 &\leq -1/2 \cdot \gamma_n \cdot t_n + 1/2 \cdot b - 1/2 \cdot \sqrt{f(0, b)} \\ &\leq -1/2 \cdot \gamma_n \cdot t_n + 1/2 \cdot b - 1/2 \cdot \sqrt{b^2} = -1/2 \gamma_n t_n \\ &< 0\end{aligned}$$

and

$$\begin{aligned}\lambda_2 &\geq -1/2 \cdot \gamma_n \cdot t_n + 1/2 \cdot b + 1/2 \cdot \sqrt{f(t_n, 0)} \\ &\geq -1/2 \cdot \gamma_n \cdot t_n + 1/2 \cdot b + 1/2 \cdot \sqrt{\gamma_n^2 \cdot t_n^2} = 1/2 \cdot b \\ &> 0.\end{aligned}$$

We also get an upper bound for $\lambda_2$ by observing that

$$f(t_n, b) \leq \gamma_n^2 t_n^2 + b^2 + 2 \cdot 0.99 b t_n < (\gamma_n t_n + b)^2$$

and hence

$$\lambda_2 < 1/2(b - \gamma_n) + 1/2(\gamma_n t_n + b) = b.$$

This shows that the positive eigenvalue is very close to zero.

We have thus shown that the trivial steady state always has one-dimensional unstable and four-dimensional stable manifold.

###### 1.1.3 System for H=0 (Normoxic region)

For $n \gg n_{cr}$ the system can be approximated by setting $H = 0$. Hence we have

$$\frac{\partial m_2}{\partial t} = D_m \nabla^2 m_2 + \kappa_{1,2} m_1 (\rho_m + \rho_p) - \kappa_{2,1} m_2 - d_2 m_2.$$

Using the equations for the time derivatives of $m_1$ and $m_2$ one easily shows that the only glioma-free steady state of the system in the normoxic region is the trivial steady state $x^*_{triv}$.

###### 1.1.4 System for H=1 (Hypoxic region)

For $n \ll n_{cr}$ the system can be approximated by setting $H = 1$. Hence we have

$$\frac{\partial m_2}{\partial t} = D_m \nabla^2 m_2 + \kappa_{1,2} m_1 (\rho_m + \rho_p) - \kappa_{2,1} m_2 + r \cdot m_2 (1 - \frac{(m_1 + m_2)}{K_m}) - d_2 m_2.$$

The unique non-trivial glioma-free steady state of the system is then given by

$$x^*_{hypo} = [0, 0, \kappa_{2,1} \tfrac{m^*_{2,hypo}}{d1}, m^*_{2,hypo}, n^*_{hypo}]$$

where

$$m^*_{2,hypo} = -\tfrac{K_m (d_2 + \kappa_{2,1} - r)}{r\left(\frac{\kappa_{2,1}}{d_1} + 1\right)}$$

$$n^*_{hypo} = -\frac{h_1 n_0 r v}{K_m d_2 h_3 + K_m h_3 \kappa_{2,1} - K_m h_3 r - h_1 r v}.$$

For the point $n^*_{hypo}$ to be in the region $n \ll n_{cr}$, assuming $r > 0$ and using our values of $n_{cr} = 0.25$ and $n_0 = v = 1$ we get the conditions

$$3h_1 < K_m h_3 \text{ and } r > -K_m h_3 \frac{d_2 + \kappa_{2,1}}{3h_1 - K_m h_3}$$

that must be satisfied to ensure $n^*_{hypo} < 0.25 = n_{cr}$. Plugging in our fixed values $K_m = 25$ and $h_1 = 3.37 \times 10^{-1}$ the first estimate evaluates to

$$h_3 > 4.044 \times 10^{-2}.$$

To analyse the stability we compute the Jacobian of the system for $H = 1$ as

$$\begin{bmatrix} \Pi\Theta\delta + \Omega t_n & (\Omega + \gamma_n)t_n - \frac{K_m \Theta t_s}{Kr} & 0 & 0 & 0 \\ -\Omega t_n & \Pi\Theta\delta - (\Omega + \gamma_n)t_n + b + \frac{K_m \Theta t_s}{Kr} & 0 & 0 & 0 \\ \Pi\Theta\kappa_{1,2} + \frac{S}{K_p} & \Pi\Theta\kappa_{1,2} + \frac{S}{K_p} & -d_1 & \kappa_{2,1} & 0 \\ -\Pi\Theta\kappa_{1,2} & -\Pi\Theta\kappa_{1,2} & \Theta & \Xi & 0 \\ \Omega h_2 & \Omega h_2 & \Omega h_3 & \Omega h_3 & \mu_5 \end{bmatrix}$$

where

$$\begin{aligned} \Omega &= \frac{h_1 n_0 r v}{K_m d_2 h_3 + K_m h_3 \kappa_{2,1} - K_m h_3 r - h_1 r v} = -n^*_{hypo}, \\ \Theta &= \frac{d_1(d_2 + \kappa_{2,1} - r)}{d_1 + \kappa_{2,1}}, \\ \Xi &= 2\Theta + r - d_2 - k21 + \frac{r\Pi\Theta}{K_m}, \\ \Pi &= \frac{K_m \kappa_{2,1}}{d_1 r}, \\ \mu_5 &= \frac{\Pi\Theta h_3 r + K_m \Theta h_3 - h_1 r v}{r}. \end{aligned}$$

This matrix has eigenvalues

$$\begin{bmatrix} \mu_1 \\ \mu_2 \\ \mu_3 \\ \mu_4 \\ \mu_5 \end{bmatrix} := \begin{bmatrix} \frac{u(r,b,t_n,t_s) + \sqrt{w(r,b,t_n,t_s)}}{2\,Kr} \\ \frac{u(r,b,t_n,t_s) - \sqrt{w(r,b,t_n,t_s)}}{2\,Kr} \\ \frac{1}{2}\,\Xi - \frac{1}{2}\,d_1 - \frac{1}{2}\,\sqrt{q} \\ \frac{1}{2}\,\Xi - \frac{1}{2}\,d_1 + \frac{1}{2}\,\sqrt{q} \\ \frac{\Pi\Theta h_3 r + K_m \Theta h_3 - h_1 r v}{r} \end{bmatrix}$$

where

$$u(r, b, t_n, t_s) := K_m \Theta t_s + (2\,K\Pi\Theta\delta + Kb)r - K\gamma_n r t_n,$$

$$w(r,b,t_n,t_s) := K^2{\gamma_n}^2 r^2 {t_n}^2 + K^2 b^2 r^2 + {K_m}^2 \Theta^2 {t_s}^2 - 2\left(2\,K^2 \Omega b + K^2 b \gamma_n\right) r^2 t_n$$
$$- 2\,(K K_m \Theta \gamma_n r t_n - K K_m \Theta b r) t_s,$$

and

$$q = (\Xi + d_1)^2 + 4\,\Theta \kappa_{2,1}.$$

**Properties of $\Theta$, $\Pi$ and $\Xi$**

First, we make some statements about the values of $\Theta$, $\Pi$, and $\Xi$.

As $r > d_2 + \kappa_{2,1}$ we know that $\Theta$ is negative. For the absolute value of $\Theta$, we have

$$|\Theta| < r \tfrac{d_1}{d_1 + \kappa_{2,1}} < r$$

and

$$|\Theta| > d_1(r - d_2 - \kappa_{2,1}) > 1/5$$

using the minimum value for $r$ and maximum value for $\kappa_{2,1}$,

so

$$1/5 < -\Theta < r. \tag{3}$$

For $\Xi$ we show that

$$-2 \cdot \sqrt{-\kappa_{2,1}\Theta} - d_1 < \Xi(r) < -d_2 \tag{4}$$

for our parameter values.

For this we note that $\Xi'(r) = -\frac{d_1}{d_1+\kappa_{2,1}}$, so $\Xi$ is monotonously decreasing in $r$.

We have $\Xi(r_{high}) = -d_2$ at

$$r_{high} = \tfrac{2d_1 d_2 + (d_1+d_2)\kappa_{2,1}}{d_1} < 2d_2 + 2\kappa_{2,1}$$

which is lower than our $r$-values. Hence since $\Xi$ is monotonously decreasing, we have $\Xi(r) < -d_2$.

We also have $\Xi(r_{low}) = -d_1 - 2 \cdot \sqrt{-\Theta\kappa_{2,1}}$ at

$$r_{low} = \frac{d_1^2 + d_1(d_2 + 2\,\kappa_{2,1}) + 2\,(d_1 + \kappa_{2,1})\sqrt{-\Theta\kappa_{2,1}}}{d_1} > d_1 + d_2 + 2\kappa_{2,1}.$$

Here we needed that $r_{low}$ is real which is true since $\Theta < 0$. We see that $r_{low}$ is greater than the $r$-values we are interested in, hence since $\Xi$ is monotonously decreasing we conclude $\Xi(r) > -2\sqrt{-\kappa_{2,1}\Theta} - d_1$.

Further, we have $\Pi > 0$.

**The term $u$ is always negative for our parameter range**

We use $t_s = 1 - t_n$, so that

$$u(r, b, t_n, t_s) = (K_m(-\Theta) - K\gamma_n r)t_n + K_m\Theta + (2\,K\Pi\Theta\delta + Kb)r.$$

We see that since $K_m(-\Theta) \overset{(3)}{<} Kr < K\gamma_n$, $u$ is a decreasing function in $t_n$.

We also have $K_m(-\Theta) \overset{(3)}{>} \frac{K_m}{5} > Kbr$ for our parameter ranges,

hence evaluating at $t_n = 0$ gives

$$u(r, b, 0, t_s) < -K_m(-\Theta) + 2K\Pi\Theta\delta r + Kbr < 0.$$

**The term $\sqrt{w}$ is always real**

We now show that $w \geq 0$ and hence $\sqrt{w}$ is always a real number. We again use the simplification $t_n = 1 - t_s$.

First, we show that $w$ is monotononously decreasing in $t_s$ for our $t_s$-values. For this we note that $w$ defines a parabola in $t_s$ with leading coefficient

$$K^2\gamma_n^2 r^2 + 2KK_m\Theta\gamma_n r + K_m^2\Theta^2.$$

Hence, since

$$K^2\gamma_n^2 r^2 + K_m^2\Theta^2 > 2KK_m\gamma_n r^2 > -2KK_m\gamma_n\Theta r.$$

the leading coefficient is greater than one and thus by (2) we need to show that $t_{s,min} > 1$.

We need to show (using $t_n = 1 - t_s$) that

$$t_{s,min} = -\frac{2\,K^2\Omega br^2 + K^2 b\gamma_n r^2 - K^2{\gamma_n}^2 r^2 + KK_m\Theta br - KK_m\Theta\gamma_n r}{K^2{\gamma_n}^2 r^2 + 2\,KK_m\Theta\gamma_n r + {K_m}^2\Theta^2} > 1.$$

Multiplying with the (positive) denominator and collecting non-negative terms becomes

$$-2\,K^2\Omega br^2 - 3\,KK_m\Theta\gamma_n r - KK_m\Theta br > {K_m}^2\Theta^2 + K^2 b\gamma_n r^2.$$

But now this is true as

$$\begin{aligned}
-2K^2\Omega br^2 &- 3KK_m\Theta\gamma_n r - KK_m\Theta br \\
&\geq -KK_m\Theta(3\gamma_n r + br) \\
&> -KK_m\Theta(3\gamma_n r) \\
&> -KK_m\Theta(r + \gamma_n r) \\
&> -KK_m\Theta(-\Theta + \gamma_n r^2) \\
&= KK_m\Theta^2 + KK_m(-\Theta)\gamma_n r^2 \\
&> K_m^2\Theta^2 + K^2 b\gamma_n r^2.
\end{aligned}$$

Hence, indeed, $t_{s,min} > 1$, so $w$ is monotonously decreasing in $t_s$ for our $t_s$-values.

At its minimum in the range $t_s \in [0,1]$, i.e. at $t_s = 1$ we have

$$w(t_s = 1) = K^2 b^2 r^2 + 2KK_m\Theta br + K_m^2\Theta^2 > 0$$

since

$$K^2 b^2 r^2 + K_m^2\Theta^2 > 4 \cdot 10^{-2} \cdot K_m^2 > K_m \cdot (4r^2 bK) > -2KK_m\Theta br.$$

Hence $w > 0$ and hence $\sqrt{w}$ is always real.

**Sign of** $\mathrm{Re}(\mu_5)$

For $\mu_5$ we know

$$\mathrm{Re}(\mu_5) = \Pi\Theta h_3 + \frac{K_m\Theta h_3}{r} - h_1 v < 0.$$

**Sign of** $\mathrm{Re}(\mu_3)$**,** $\mathrm{Re}(\mu_4)$

Clearly $\mathrm{Re}(\mu_3) \leq \mathrm{Re}(\mu_4)$. Our goal is show $\mathrm{Re}(\mu_4) < 0$. From the lower bound for $\Xi$ in (4) we already know that $\sqrt{q}$ is a real number.

Now we can infer that

$$\mathrm{Re}(\mu_3) \leq \mathrm{Re}(\mu_4) = \tfrac{1}{2}(\Xi - d_1) + \sqrt{q} \overset{\Theta<0}{\leq} \tfrac{1}{2}(\Xi - d_1) + \tfrac{1}{2}\sqrt{(\Xi + d_1)^2} \overset{(3)}{=} \Xi < 0.$$

**Sign of** $\mathrm{Re}(\mu_2)$

For $\mu_2$ we know that $u < 0$ and hence surely $\mathrm{Re}(\mu_2) < 0$.

**Sign of** $\mathrm{Re}(\mu_1)$

For $\mu_1$ we have that

$$\begin{aligned} u^2 - w = {} & 4\,KK_m\Pi\Theta^2\delta r t_s \\ & + 4\left(K^2\Pi^2\Theta^2\delta^2 + K^2\Pi\Theta b\delta - \left(K^2\Pi\Theta\delta\gamma_n - K^2\Omega b\right)t_n\right)r^2. \end{aligned}$$

Using $t_n = 1 - t_s$ and solving for $b$ yields

$$b_{crit} = \frac{\left(K\Pi^2\Theta^2\delta^2 - K\Pi\Theta\delta\gamma_n\right)r + \left(K\Pi\Theta\delta\gamma_n r + K_m\Pi\Theta^2\delta\right)t_s}{K\Omega r t_s - (K\Pi\Theta\delta + K\Omega)r} \tag{5}$$

At this point we have $\mu_1 = 0$, hence the stability changes.

We can further compute

$$\frac{\partial}{\partial b}(u^2 - w)(b_{crit}) = 4K^2r^2(\Pi\Theta\delta + \Omega t_n) < 0$$

since $\Theta, \Omega < 0$.

Hence at $b_{crit}$ the system changes from an unstable to a stable state.

**Summary for H=1**

To summarize everything we have shown in this subsection that given

$$3h_1 < K_m h_3$$

and

$$r > -K_m h_3 \tfrac{d_2+\kappa_{2,1}}{3h_1 - K_m h_3}$$

we can approximate a glioma-cell free steady state of our system by

$$x^*_{hypo} = [0, 0, \kappa_{2,1}\tfrac{m^*_{2,hypo}}{d_1}, m^*_{2,hypo}, n^*_{hypo}]$$

where

$$m^*_{2,hypo} = -\tfrac{K_m(d_2+\kappa_{2,1}-r)}{r\left(\frac{\kappa_{2,1}}{d_1}+1\right)}$$

and

$$n^*_{hypo} = -\tfrac{h_1 n_0 r v}{K_m d_2 h_3 + K_m h_3 \kappa_{2,1} - K_m h_3 r - h_1 r v}.$$

The linear stability for our parameter values can be characterised as follows:

All eigenvalues are real and we have $\mu_2, \mu_3, \mu_4, \mu_5 < 0$, so the steady state has at least 4-dimensional stable manifold.

If

$$b > \frac{\left(K\Pi^2\Theta^2\delta^2 - K\Pi\Theta\delta\gamma_n\right)r + \left(K\Pi\Theta\delta\gamma_n r + K_m\Pi\Theta^2\delta\right)t_s}{K\Omega r t_s - (K\Pi\Theta\delta + K\Omega)r}$$

we have 5-dimensional stable manifold and if

$$b < \frac{\left(K\Pi^2\Theta^2\delta^2 - K\Pi\Theta\delta\gamma_n\right)r + \left(K\Pi\Theta\delta\gamma_n r + K_m\Pi\Theta^2\delta\right)t_s}{K\Omega r t_s - (K\Pi\Theta\delta + K\Omega)r}$$

we have 4-dimensional stable manifold and 1-dimensional unstable manifold.

##### 1.1.5 Conclusion of the Stability Analysis

In our stability analysis for the model proposed in the main text we analysed the behaviour of the diffusion-free system around its glioma-free steady states. Our analysis shows that for our parameter values the trivial steady state always has one-dimensional unstable and four-dimensional stable manifold. This state is the only macrophage-free state of the system. Hence to be macrophage-free the system also requires the absence of tumour cells.

In the hypoxic region, i.e. $n \ll n_{cr}$ we saw that there is a bifurcation occurring. We wrote down the dependence on the value of the glioma proliferation rate $b$ which emphasizes that the glioma proliferation rate is a critical parameter that determines whether the tumour will die out eventually. However the expression (5) is also dependent of all parameters except the saturation rate $K_p$, the recruitment rate $S$, the glioma cell oxygen consumption rate $h_2$ and the $M1$ to $M2$ switch rate $\kappa_{1,2}$. All other parameters have an impact on the stability of the steady state in the hypoxic region.

#### 1.2 Random sampling of parameters for the sensitivity analysis

For the sampling, we consider using a Latin hypercube sampling (LHS). This method is particularly effective for sampling high-dimensional spaces because it ensures that the entire range of each variable is explored uniformly. Unlike simple random sampling, which might cluster samples in some regions while leaving others underrepresented, LHS stratifies the sampling space, leading to a more efficient and comprehensive exploration of the parameter space [20]. This stratification can be especially beneficial in the context of our 5D nonlinear system, where understanding the influence of multiple interacting variables is crucial. In LHS, the range of each input parameter is divided into equal probability intervals, and samples are drawn such that each interval is sampled exactly once. This helps in obtaining a better approximation of the underlying distribution and reduces the variance of the estimation. For the majority of our parameters, we determined that the uniform probability distribution was the most suitable for describing the ranges of possible values. This distribution is appropriate when we lack an estimate of a most likely value for a parameter. However, for three specific parameters—growth rate, infiltration rate, and glioma consumption rate—we utilized a triangular distribution based on the method described by Massey (2016) [21]. This choice was due to our ability to identify maximum and minimum values, as well as to estimate a most likely value, despite lacking further information about the distribution's shape. If we acquire more comprehensive data about the potential values of the parameters, other distributions might prove to be more appropriate.

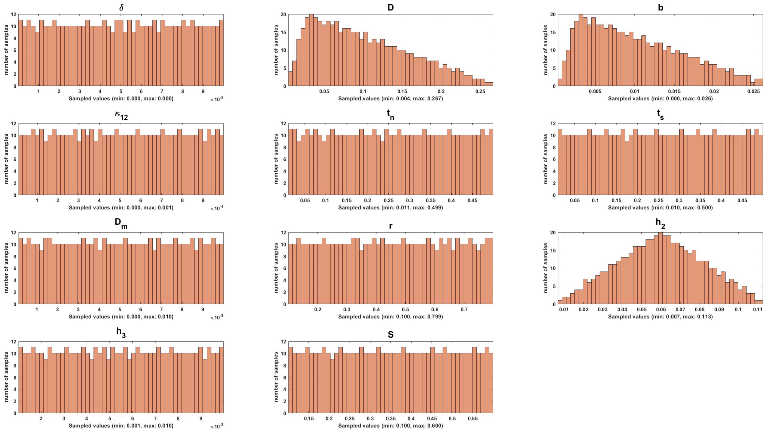


**Fig. 1** Histograms of the parameter ranges used in the PRCC sensitivity analysis.

##### 1.3 Choosing the best machine learning model for investigating the further result

In addition to the data preprocessing, we analyze the dataset after the standardization (using Z-score) to explore it via a wide variety of models. We first consider the standardized dataset into train and test data (80% training and 20% validation set) and then implement the train and validation set to the model to figure out the MSE of each model and the possible overfitting. The bar plots in Fig . 10, illustrate the MSE scores for ten different regression models on both training and test datasets. Gradient Boosting is selected as the optimal model due to its relatively low and closely matched MSE scores between the training (7.8764 months$^2$) and test datasets (8.5879 months$^2$), indicating robust performance with minimal overfitting. This is contrasted with models like Decision Tree, which shows zero MSE in training but significantly higher MSE in testing, suggesting overfitting. The bar plots effectively highlight the differences in model performance, guiding the selection of the most stable and generalizable model for further analysis.

### References

[1] R. Seydel, Practical Bifurcation and Stability Analysis, Vol. 5 of Interdisciplinary Applied Mathematics, Springer New York, 2010. doi:10.1007/978-1-4419-1740-9.
URL `https://link.springer.com/10.1007/978-1-4419-1740-9`

[2] K. R. Swanson, R. C. Rockne, J. Claridge, M. A. Chaplain, E. C. Alvord Jr, A. R. Anderson, Quantifying the role of angiogenesis in malignant progres-

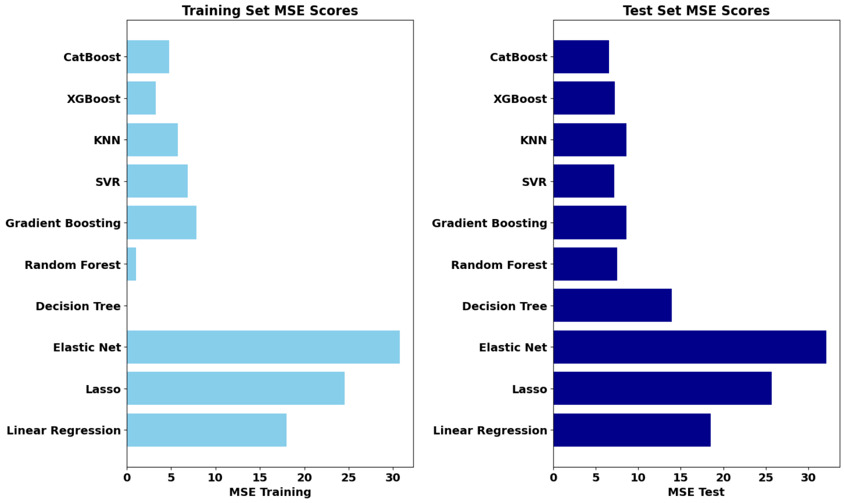


**Fig. 2** Comparison of Mean Squared Error (MSE) Across Various Regression Models for Training and Test Datasets to select the best model based on the MSE values and minimum difference between test and train cases.

sion of gliomas: in silico modeling integrates imaging and histology, Cancer research 71 (24) (2011) 7366–7375.

[3] H. L. Harpold, E. C. Alvord Jr, K. R. Swanson, The evolution of mathematical modeling of glioma proliferation and invasion, Journal of Neuropathology & Experimental Neurology 66 (1) (2007) 1–9.

[4] M. Badoual, C. Gerin, C. Deroulers, B. Grammaticos, J.-F. Llitjos, C. Oppenheim, P. Varlet, J. Pallud, Oedema-based model for diffuse low-grade gliomas: application to clinical cases under radiotherapy, Cell proliferation 47 (4) (2014) 369–380.

[5] S. E. Eikenberry, T. Sankar, M. C. Preul, E. J. Kostelich, C. Thalhauser, Y. Kuang, Virtual glioblastoma: growth, migration and treatment in a three-dimensional mathematical model, Cell proliferation 42 (4) (2009) 511–528.

[6] J. McDaniel, E. Kostelich, Y. Kuang, J. Nagy, M. C. Preul, N. Z. Moore, N. L. Matirosyan, Data assimilation in brain tumor models, Mathematical methods and models in biomedicine (2013) 233–262.

[7] A. Buonfiglioli, D. Hambardzumyan, Macrophages and microglia: the cerberus of glioblastoma, Acta neuropathologica communications 9 (1) (2021) 1–21.

[8] R. Eftimie, C. Barelle, Mathematical investigation of innate immune responses to lung cancer: The role of macrophages with mixed phenotypes,

Journal of Theoretical Biology 524 (2021) 110739.

[9] W. E. Hoffman, F. T. Charbel, G. Edelman, K. Hannigan, J. I. Ausman, Brain tissue oxygen pressure, carbon dioxide pressure and ph during ischemia, Neurological research 18 (1) (1996) 54–56.

[10] A. Carreau, B. E. Hafny-Rahbi, A. Matejuk, C. Grillon, C. Kieda, Why is the partial oxygen pressure of human tissues a crucial parameter? small molecules and hypoxia, Journal of cellular and molecular medicine 15 (6) (2011) 1239–1253.

[11] I. Stamper, M. Owen, P. Maini, H. Byrne, Oscillatory dynamics in a model of vascular tumour growth-implications for chemotherapy, Biology direct 5 (1) (2010) 1–17.

[12] A. Matzavinos, C.-Y. Kao, J. E. F. Green, A. Sutradhar, M. Miller, A. Friedman, Modeling oxygen transport in surgical tissue transfer, Proceedings of the National Academy of Sciences 106 (29) (2009) 12091–12096.

[13] G. Powathil, M. Kohandel, M. Milosevic, S. Sivaloganathan, et al., Modeling the spatial distribution of chronic tumor hypoxia: implications for experimental and clinical studies, Computational and mathematical methods in medicine 2012 (2012).

[14] C. D. Eggleton, T. K. Roy, A. S. Popel, Predictions of capillary oxygen transport in the presence of fluorocarbon additives, American Journal of Physiology-Heart and Circulatory Physiology 275 (6) (1998) H2250–H2257.

[15] D. Goldman, A. S. Popel, A computational study of the effect of capillary network anastomoses and tortuosity on oxygen transport, Journal of theoretical biology 206 (2) (2000) 181–194.

[16] C. J. Kelly, M. Brady, A model to simulate tumour oxygenation and dynamic [18f]-fmiso pet data, Physics in Medicine & Biology 51 (22) (2006) 5859.

[17] D. R. Grimes, C. Kelly, K. Bloch, M. Partridge, A method for estimating the oxygen consumption rate in multicellular tumour spheroids, Journal of The Royal Society Interface 11 (92) (2014) 20131124.

[18] P. Vaupel, F. Kallinowski, P. Okunieff, Blood flow, oxygen and nutrient supply, and metabolic microenvironment of human tumors: a review, Cancer research 49 (23) (1989) 6449–6465.

[19] The Sage Developers, SageMath, the Sage Mathematics Software System (Version 10.3), `https://www.sagemath.org` (2024).

[20] M. D. McKay, R. J. Beckman, W. J. Conover, A comparison of three methods for selecting values of input variables in the analysis of output from a computer code, Technometrics 42 (1) (2000) 55–61.

[21] S. C. Massey, Multi-scale modeling of paracrine pdgf-driven glioma growth and invasion, Ph.D. thesis (2016).
